# Sterol 3-beta-Glucosyltransferase TRANSPARENT TESTA15 Controls Seed Development and Flavanol Accumulation through its Role in Vacuole Biogenesis and Maintenance in Arabidopsis

**DOI:** 10.1101/2023.09.12.557332

**Authors:** Elodie Akary, Adeline Berger, François Perreau, Anne Frey, Alexandra To, Sylvie Citerne, Hubert Schaller, Samantha Vernhettes, Olivier Grandjean, Nathalie Nesi, Annie Marion-Poll, Loïc Lepiniec, Isabelle Debeaujon

## Abstract

The Arabidopsis sterol 3-beta-glucosyltransferase UGT80B1/TRANSPARENT TESTA15 (TT15) catalyzes sterol glucoside biosynthesis. Its loss of function causes reduced seed size, defective flavanol, polysaccharide and lipid polyester deposition at the seed coat and reduced seed dormancy. How TT15 controls seed development and physiology is unknown. Here we show that *tt15* mutants exhibit seed lethality with incomplete penetrance and maternal determinism that is correlated with endosperm cellularization defects, together with an increased sensitivity of seed germination to exogenous abscisic acid and paclobutrazol. We also reveal that flavanol deposition in the vacuole during *tt15* seed development triggers premature endothelium cell death. An autoimmune-like syndrome characterized by callose and H_2_O_2_ accumulation was detected in endothelium at the seed abaxial pole. Similar phenotypes were observed with *tt9/gfs9,* a mutant defective in endomembrane trafficking and homotypic vacuole fusion. Double mutant analysis showed that *tt9* partially rescued *tt15* endothelium phenotypes. Consistent with seed mutant phenotypes, *TT15* promoter activity was detected in endothelium and endosperm and TT15 protein was located mainly at the vacuolar membrane (tonoplast). Using fluorescence recovery after photobleaching, we demonstrated that tonoplast fluidity was increased in *tt15* roots. Altogether our data suggest that TT15 regulates seed development and flavanol accumulation by modulating vacuole biogenesis and maintenance.

## INTRODUCTION

Seed development in Angiosperms is initiated by double fertilization, leading to the formation of a diploid embryo and a triploid endosperm. The two siblings develop concomitantly within the surrounding maternal tissue or seed coat (also called testa), which is derived from the post-fertilization differentiation of the two ovule integuments (inner integument or ii, and outer integument or oi) in Arabidopsis. When morphogenesis is completed, the embryo grows at the expense of the nurturing endosperm. The three seed components need to exchange signals to ensure coordinated development, maturation and differentiation that determine final seed traits, among which auxin and sugars play prominent roles (Ingram, 2010; Figueiredo and Köhler, 2016; Robert, 2019). Seed germination begins with the uptake of water during imbibition of quiescent dry seeds and ends up when hypocotyl expansion triggers protrusion of the embryo radicle through the seed envelopes. These ones consist in a dead brown seed coat and a single layer of live endosperm in Arabidopsis. A dormant seed is unable to germinate, even in favourable environmental conditions. The control of germination results from the competitive interaction between embryonic growth potential and mechanical restraint imposed by surrounding tissues. Abscisic acid (ABA) and gibberellins (GAs) are diterpenoid hormones (Supplemental Figure S1) acting antagonistically in seed dormancy and germination control. The ABA/GA balance is an integrator of environmental and metabolic clues favourable to seed germination such as water, oxygen, temperature, light and nitrate (North et al., 2010).

Sterols, a class of lipids of terpenic origin (Supplemental Figure S1), play crucial roles in plant development and growth as components of membranes and as precursors for steroidal hormones brassinosteroids (BR) and steroidal specialized metabolites. They are also essential for proper seed development and physiology (Schaller, 2004; Mamode Cassim et al., 2019; Shimada et al., 2021). In most plants and fungi, some animals and a few bacteria, sterols are present not only as free sterols (FS) but also conjugated as steryl glycosides (SG) and acyl steryl glycosides (ASG). The sugar moiety (generally a D-glucose) is attached to the 3β-hydroxy group at the C3-atom of a sterol. It increases the size of the hydrophilic head-group of the lipid and thus changes its biophysical properties. The conversion of membrane-bound FS to SG is catalyzed by nucleoside diphosphate (NDP)-sugar-dependent sterol glycosyltransferases (Grille et al., 2010). Sterol glycosyltransferases (EC 2.4.1.173) play important roles in plant metabolic plasticity during adaptive responses (Grille et al., 2010; Ferrer et al., 2017). In Arabidopsis, two uridine diphosphate (UDP)-glucose:sterol glucosyltransferases have been identified, namely UGT80A2 and UGT80B1, and their biological functions explored by a reverse genetics approach (Warnecke et al., 1997; DeBolt et al., 2009). Phenotypic analysis of the corresponding single mutants revealed important perturbations in seed development specifically for *ugt80b1*, namely a *transparent testa (tt)* phenotype (pale seeds compared to brown wild-type seeds), a reduced seed size, a loss of cutin and suberin at the seed coat, but no impact on cellulose biosynthesis in vegetative parts (DeBolt et al., 2009). The *ugt80b1* mutant appeared to be allelic to the *transparent testa15* (*tt15)* mutant previously identified by Focks et al. (1999). Recently, specific roles for both enzymes in polysaccharide accumulation at the level of seed coat epidermal cells (SCE or oi2 cells) were inferred from a thorough cytological, chemical and physico-chemical characterization of *ugt80A2* and *ugt80B1/tt15* mutants. This study revealed that *tt15* oi2 cells do not release properly their mucilage due to a localized increase in the deposition of secondary cell wall polymers at the level of radial oi2 cell walls (Berger et al., 2021). Albeit they are classified in the family 1 of plant UDP glycosyltransferases (UGT), Arabidopsis UGT80B1/TT15 and its paralog UGT80A2 are very divergent from other plant UGTs because they do not contain an obvious Plant Secondary Product Glycosyltransferase motif (PSPG) and are more closely related to non-plant UGT families (Caputi et al., 2012). The sequence homology between UGT80B1/TT15 and its orthologs is restricted to the catalytic region including the Putative Steroid-Binding Domain (PSBD) (Warnecke et al., 1999) and the C-terminal PROSITE consensus sequence for family 1 glycosyltransferases (UGT). *In vitro* enzyme assays and the analysis of SG in seeds of single mutants revealed discrepancies between both paralogs. Indeed, if UGT80A2 is responsible for the bulk production of SGs, UGT80B1/TT15 is involved in the production of minor but probably critical SGs, such as campesteryl and brassicasteryl glucosides (Stucky et al., 2015). Another meaningful difference between both paralogs is their subcellular localization as determined by proteomics analyses. UGT80B1/TT15 was detected at the vacuolar membrane or tonoplast (Carter et al., 2004; Jaquinod et al., 2007) and UGT80A2 at the plasma membrane (Marmagne et al., 2007; Zhang and Peck, 2011). On the other hand, both paralogs were shown to be peripheral membrane proteins, consistent with the absence of transmembrane domains (Ramirez-Estrada et al., 2017).

Seed coat colour in Arabidopsis is conferred by flavanols, a subclass of flavonoids involving proanthocyanidins (also called condensed tannins) and their flavan-3-ol monomers. Arabidopsis wild-type seeds synthesize exclusively procyanidins (PC) resulting from the condensation of epicatechin (EC) monomers (Routaboul et al., 2006) (Supplemental Figure S2). PC and EC accumulate as colourless compounds in vacuoles of tannin-producing cells, namely the endothelium (ii1 cells), the micropylar region (a few ii1’ cells) and the chalazal pigment strand of the seed coat and are further oxidized as brown pigments by the TT10 laccase upon seed desiccation (Debeaujon et al., 2003; Pourcel et al., 2005). Mutants with altered seed coat colour, thus defective in PC metabolism (*tt*; *tt glabra* or *ttg*; *tannin-deficient seeds* or *tds*; *banyuls* or *ban*) were instrumental in establishing many steps of the flavonoid biosynthetic pathway (Koornneef, 1990; Lepiniec et al., 2006). PC production is triggered by ovule fertilization and ends around the heart stage of embryo development (Debeaujon et al., 2003; Figueiredo and Köhler, 2014). It is tightly regulated spatio-temporally by a complex of transcriptional regulators involving a Myeloblastosis (MYB), a basic Helix-Loop-Helix (bHLH) and a WD-repeat (WDR) protein (MBW complex) encoded by the *TT2*, *TT8* and *TTG1* genes, respectively (Lepiniec et al., 2006). TTG2, a WRKY-type transcription factor, was proposed to regulate vacuolar transport steps (Johnson et al., 2002; Gonzalez et al., 2016). Upstream the MBW complex, at least two other transcription factors, namely a WIP-type zinc finger and a MADS encoded by the *TT1* and *TT16* genes respectively, control endothelium identity and thus its competency to accumulate flavanols (Nesi et al., 2002; Sagasser et al., 2002). Proanthocyanidins have substantial antioxidant activity, and the ability to chelate metals and to cross-link with proteins and cell wall polysaccharides. These specialized metabolites reinforce coat-imposed dormancy and seed longevity (Debeaujon et al., 2000). Moreover, their role in the regulation of seed size and the control of post-zygotic reproductive barriers is questioned (Garcia et al., 2005; Dilkes et al., 2008; Doughty et al., 2014; Batista et al., 2019; Köhler et al., 2021). The mechanisms mediating flavanol transport to the vacuole in wild-type seeds also are still a matter of debate (Dixon and Sarnala, 2020). The current working model for flavanol trafficking based on Arabidopsis and Medicago biochemistry and genetics postulates that after biosynthesis by a metabolon anchored at the external side of the ER, EC is glycosylated and transported to the vacuole where it would hypothetically be hydrolyzed by a glycosidase and polymerized to PCs before migrating to the cell wall according to an unknown mechanism (Winkel, 2019; Dixon and Sarnala, 2020) (Supplemental Figure S2). The vacuolar transport of glycosylated EC involves the tonoplastic Multidrug And Toxin Extrusion (MATE) transporter TT12 / Detoxification 41 (DTX41) / TDS3 (Debeaujon et al., 2001; Marinova et al., 2007; Zhao and Dixon, 2009; Appelhagen et al., 2014) that would be energized by the tonoplastic P_3A_-ATPase TT13/Autoinhibited H^+^-ATPase isoform 10 (AHA10) / TDS5 (Baxter et al., 2005; Appelhagen et al., 2014; Appelhagen et al., 2015). Another actor is the glutathione S-transferase (GST) TT19/GST26/GSTF12 that may work as a ligandin to protect the flavonoid molecule from oxidative degradation in the cytosol until it reaches its dedicated transporter at the tonoplast (Kitamura et al., 2010). Vesicle trafficking also participates in vacuolar transport of flavanols in Arabidopsis seed coats. The peripheral membrane protein TT9/GREEN FLUORESCENT SEED9 (GFS9) localized at the Golgi is involved in vacuole biogenesis and genetically interacts with the trans-Golgi network (TGN)-located ECHIDNA (ECH) protein (Ichino et al., 2014; Ichino et al., 2020). The Arabidopsis AAA ATPase VACUOLAR PROTEIN SORTING4/SUPPRESSOR OF K^+^ TRANSPORT GROWTH DEFECT1 (VPS4/SKD1) is a subunit of the endosomal sorting complexes required for transport (ESCRT) machinery involved in the formation of multivesicular bodies (MVB). Seeds expressing a dominant-negative version of AtSKD1 have a *tt* phenotype and mucilage defects (Shahriari et al., 2010a). Flavanol-accumulating cells in the endothelium of various *tt/tds* mutants including *tt15* exhibit defects in biogenesis of the central vacuole, suggesting a link between flavanol accumulation and vacuole morphology (Debeaujon et al., 2001; Abrahams et al., 2003; Baxter et al., 2005; Kitamura et al., 2010; Appelhagen et al., 2014). Anthocyanin flavonoid pigment sequestered in ER-derived vesicle-like structures was shown to be targeted directly to the protein storage vacuole in a Golgi-independent manner in Arabidopsis seedlings (Poustka et al., 2007). On the same line, several works demonstrated that autophagy mechanisms have a role in anthocyanin transport to the vacuole (Külich and Zarsky, 2014; Chanoca et al., 2015). Whether these anthocyanin trafficking routes are also used by flavanols remains to be investigated (Bassham, 2015; Chanoca et al., 2015).

The molecular mechanisms by which TT15 and SG regulate seed development and flavanol accumulation in the seed coat are still unclear. Here, we show that *TT15* disruption causes seed lethality with variable penetrance and maternal determinism that is correlated with impaired endosperm cellularization. We also reveal that vacuole biogenesis in endothelial cells is affected, leading to their premature degenerescence and death and consequently to a *tt* phenotype which is associated with a polarized accumulation of radical oxygen species (ROS) and callose at the curving zone. Premature endothelium cell death (PECD) is suppressed in the absence of flavanols. The TT15 protein was located mainly at the vacuolar membrane or tonoplast and demonstrated to decrease its fluidity. Consistent with these findings, TT15 was shown to genetically interact with TT9, a protein involved in vacuole biogenesis. We propose that TT15 and SG are required for the modulation of vacuole functions required for proper endothelium-endosperm crosstalk and flavanol deposition. As a consequence, they reinforce seed dormancy by increasing seed coat impermeability properties. We also discuss the potential roles of TT15 and SG as links between vacuole, flavonoids and seed development in the establishment of post-zygotic reproductive barriers.

## RESULTS

### The *tt15* Mutations Cause Seed Lethality and Seedling Developmental Defects with Incomplete Penetrance

Previous studies on the characterization of *tt15* mutants have identified the perturbation of several important seed traits and established the pleiotropic complexity of the mutations using only one allele either in Columbia (Col) (Focks et al., 1999; Stucky et al., 2015) or Wassilewskija (Ws) background (DeBolt et al., 2009; Routaboul et al., 2012). To progress further in our understanding of TT15 functions, we started our work with an allelic series of three alleles in Col-0 and three alleles in Ws-4 from the Versailles T-DNA collection (Supplemental Table S1). The nature and position of the mutations are shown in Supplemental Figure S3A. The alleles harbour a similar pale grayish brown seed coat colour (Supplemental Figure S3B). Sterol profiling of the *tt15-2* allele in Ws-4 background showed a reduction in SG and ASG (Supplemental Figure S3C), as expected for a mutant affected in a sterol glucosyl transferase and similarly to the previously characterized *tt15* alleles by DeBolt et al. (2009) and Stucky et al. (2015).

Reciprocal crosses between *tt15-2* and wild type Ws-4 (Table 1) showed that the seed colour phenotype is maternally inherited, which is consistent with the fact that the seed coat originates from the ovule integuments. They also revealed that the *tt15-2* mutation, which behaves as a recessive trait, exhibits a slightly reduced transmission through the female parent. Indeed the number of F2 plants producing a *tt15* phenotype was lower than the number expected for Mendelian inheritance of a recessive mutation (3:1 ratio *TT15*:*tt15* seeds).

**Table 1.**
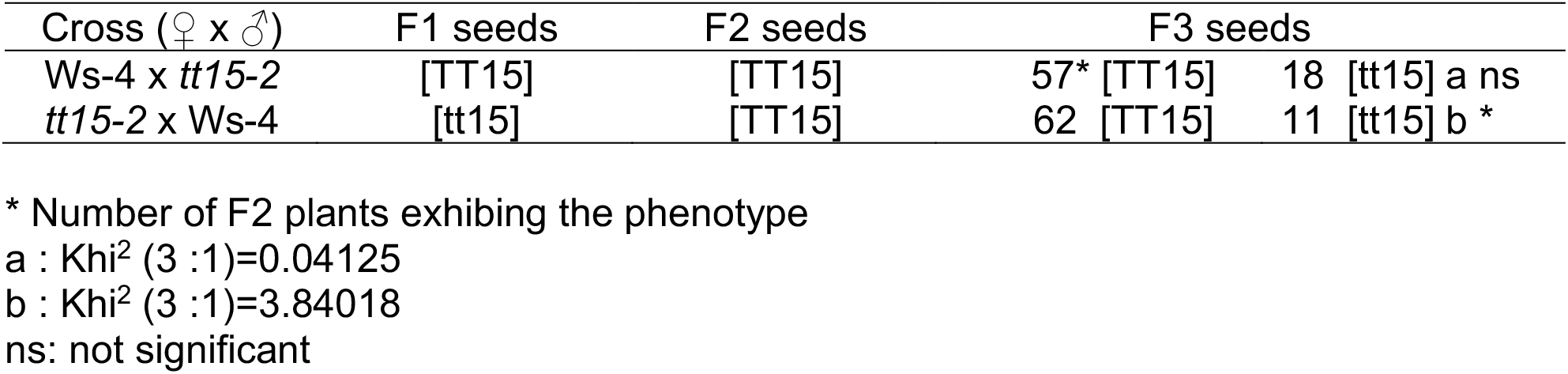
Genetic Determinism and Transmission of the *tt15-2* Seed Coat Colour.

Systematic observations of *tt15-2* seed batches below binoculars directly after harvest without cleaning (Figure 1, A and B) revealed four main phenotypic classes based on embryo development (Figure 1C). Observations done on a bulk of 3801 seeds harvested from three independent *tt15-2* plants established that around 88.5% seeds resembled wild-type seeds (class I), 8% were smaller seeds at the cotyledonary stage (class II), 0.8% seeds were mostly at the walking stick stage (class III), and 1.8% were aborted seeds with precocious arrest of embryo development at globular to torpedo stages (class IV). All seeds were able to germinate, except the ones from classes III and IV. Additionally a few seeds from class I (around 0.5% from total seeds) revealed extreme testa weakness at their abaxial pole, letting the embryo partially exit in the course of their development and growth in the silique (Figure 1D). These observations led us to conclude that embryo abortion in *tt15-2* exhibits an incomplete penetrance. Because variable frequencies of seed abortion were frequently observed in the greenhouse upon challenging growth conditions, seed lethality (aborted seed number) was quantified with seed progenies from plants grown in optimal environmental conditions in a growth chamber (Figure 1, E and F). Twenty five siliques (five siliques from five plants) developed from tagged flowers were harvested at 15 daf for *tt15-2*, *tt15-9* and their corresponding wild types Ws-4 and Col-0, respectively. Silique clearing enabled to distinguish aborted seeds as brown and flattened envelopes (Figure 1E, arrowheads). Aborted ovules appearing as white dried structures were not observed in most siliques. Differences in levels of chlorophyll breakdown between WT and mutants and between both WTs were also observed, probably revealing discrepancies in the timing of seed maturation. The mean percentage of seed lethality was significantly higher in mutants compared with corresponding WTs. Abortion was also higher in mutant in Col-0 background than in mutant in Ws-4 background (Figure 1F). Abortion scores per silique oscillated between 0% and 36.7% for *tt15-2* and between 10% and 60% for *tt15-9* compared with 0% to 5.4% in Ws-4 and 0% to 16.1% in Col-0 (Supplemental Table S3). To determine which seed compartment controls embryo abortion in a *tt15* mutant background, we performed reciprocal crosses between *tt15* and corresponding wild types. Our data showed that F1 seed lethality caused by *tt15* is maternally determined (Figure 1G; Supplemental Table S3).

**Figure 1.**
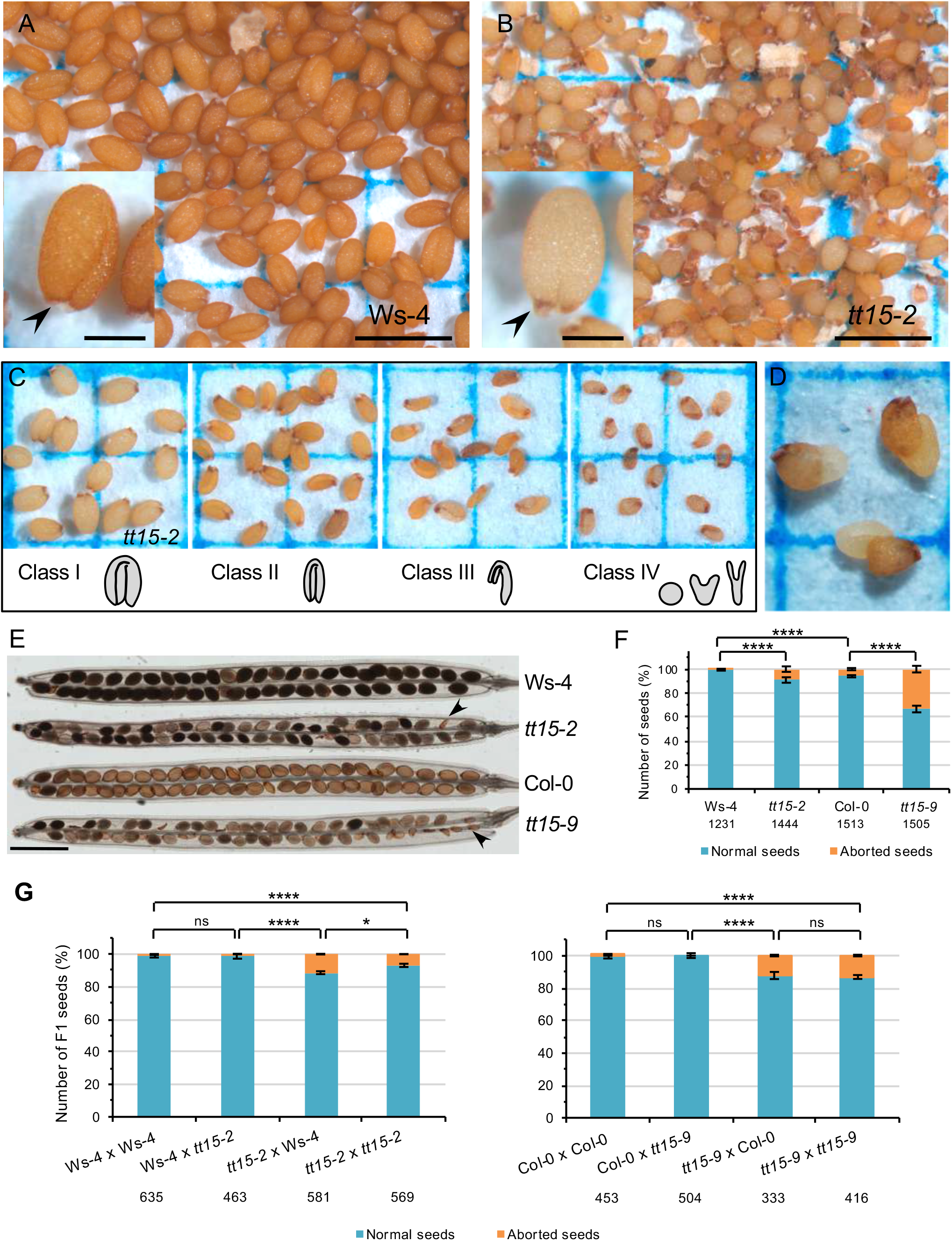
*TT15* Disruption Causes Seed Lethality and Seed Coat Developmental Defects with Incomplete Penetrance and Maternal Inheritance. **(A-B)** Mature seed phenotypes showing that *tt15* mutant seeds **(B)** are paler, being pigmented only at the micropyle-chalaza pole (arrowhead in inset) and are smaller than wild-type seeds **(A)**. Abnormal seed shapes (from shriveled to aborted seeds) are also observed in **(B)**. A normally shaped seed is shown in inset. **(C)** Different seed phenotypic classes encountered in a *tt15-2* plant progeny. Schemes refer to prevalent embryo stages observed in mature seeds. **(D)** Dry mature seeds of *tt15-2* with embryos emerging from the testa at the level of the curving zone. **(E)** and **(F)** Observation of seed lethality in maturing siliques. **(E)** Cleared siliques. Whole-mounts at 15 days after flowering are shown. Arrowheads point to aborted seeds. **(F)** Quantification of aborted seeds realized on 25 siliques per genotype (5 siliques from 5 plants). Sample size is indicated below each bar. Error bars represent standard errors (n=25). **(G)** Classification of F1 seeds derived from reciprocal crosses between wild types and corresponding *tt15* mutants based on the seed lethality phenotype. Sample size is indicated below each bar. ns, not significant. Error bars represent standard errors (n=13 for Ws-4 serie and n=8 for Col-0 serie, with n being the number of analyzed siliques). Asterisks in **(F)** and **(G)** indicate significant differences for aborted seeds using the non-parametric Mann-Whitney U test (****α=0.1% ***α=1%; ** α=2.5%; * α=5%; ns, not significant at α=5%). Bar = 1 mm (250 μm in insets) in **(A)** and **(B)** and 1.5 mm in **(E)**.

The pattern of seed coat pigmentation in *tt15-2*, with its characteristic dark brown chalaza-micropyle area and pale seed body (Figure 1B, inset; Supplemental Figure S4A) is similar to the ones of *tt1-4*, *tt9-1* and *tt16-1* mutants (Supplemental Figure S4, B-D, insets). Intriguingly, these three mutants also exhibit some seed abortion. DeBolt et al. (2009) have shown that *ugt80B1*, a *tt15* allele from the University of Wisconsin T-DNA collection (Sussman et al., 2000) exhibited a reduced seed weight. We confirmed these results with our alleles and showed that weight reduction was consistent with a smaller seed size as expected, but without modification in seed shape (Supplemental Figure S5).

Other phenotypes expressed with incomplete penetrance concern seedling development. If most mutant seedlings in a plant progeny harbor a wild-type phenotype (Supplemental Figure S6, A-C), a small fraction (between 1 to 2%) exhibits various developmental defects such as an aberrant cotyledon number (tricotyly, monocotyly, no cotyledons; Supplemental Figure S6, D, E and I, respectively), white cotyledon tips (Supplemental Figure S6F), root-like structures at cotyledon tips (Supplemental Figure S6G), transformation of a cauline apex into a root-like apex (Supplemental Figure S6H). Atypical adhesion of the aleurone layer (peripheral endosperm) and seed coat to the root tip and an irregular epidermal surface were also observed (Supplemental Figure S6, H and I). Such phenotypes could be observed with all alleles, especially in seed classes II and III.

### Sensitivity of Seed Germination to Exogenous Abscisic Acid and Paclobutrazol is Increased in *tt15* Mutant Backgrounds

A collection of *tt* mutants affected in flavanol metabolism in the seed coat was previously shown to exhibit a reduced primary dormancy, positively correlated with an increased testa permeability to tetrazolium salts (Koornneef, 1981; Debeaujon et al., 2000). Here we could extend these observations to *tt15* mutant seeds and explored further the germination phenotype using a *tt15* allelic series in Ws-4 and Col-0 backgrounds. To avoid a bias due to the seed lethality phenotype (see above), seed lots were cleaned before use to remove the dead seed fractions corresponding to classes III and IV. Freshly harvested seeds from the *tt15* alleles in Ws-4 background exhibited a reduced primary dormancy. For *tt15* alleles in Col-0 background, differences in dormancy were less significant than in Ws-4 (Supplemental Figure S7A). Because a *tt7* mutant affected in the production of dihydroquercetin derivatives (Supplemental Figure S2) was previously demonstrated to be resistant to thermoinhibition (Tamura et al., 2006), we were tempted to investigate *tt15* seed germination tolerance to high temperature. We used a temperature of 34°C previously shown to completely inhibit Arabidopsis wild-type seed germination (Tamura et al., 2006), compared to a control temperature of 22°C. However the observed differences were not statistically significant (Supplemental Figure S7B). On the other hand, seed coat permeability to tetrazolium salts was increased similarly in all alleles compared with their wild types (Supplemental Figure S7C).

To determine whether *tt15* reduced seed dormancy may also have an hormonal component residing in endosperm and/or embryo beside the seed coat physicochemical defects, we assessed *tt15* seed germination response to increasing doses of the dormancy inducer and germination inhibitor ABA after stratification. Germination *sensu stricto* of *tt15* seeds estimated by the percentage of seeds exhibiting radicle protrusion through the seed envelopes was clearly more sensitive to ABA than the corresponding wild types in both accession backgrounds (Figure 2A). Cotyledon greening during photoautotrophic seedling establishment was affected only with the alleles in Ws-4 background and not with the ones in Col-0 (Figure 2B). However hypersensitivity of cotyledon greening to the GA biosynthesis inhibitor paclobutrazol (PAC) was more contrasted between mutants and their wild types, and shared by both accession backgrounds (Figure 2C), possibly pointing to a chloroplast biogenesis defect involving GA in *tt15*, independently from the ABA-related component influencing germination *sensu stricto*. Dry Ws-4 seeds exhibited more ABA than Col-0 seeds, which may explain the differential seed dormancy and germination behaviours of Ws-4 and Col-0. On the other hand, *tt15* mutant seeds did not exhibit any reduction in ABA compared with their corresponding wild types (Supplemental Figure S7D). Hypersensitivity to exogenous ABA would explain the increased need for GA to germinate suggested by hypersensitivity to PAC. Thus in *tt15* mutant seeds either GA biosynthesis and/or ABA catabolism upon imbibition may be affected, increasing the ABA/GA ratio, or ABA and/or GA signalling are impacted. Measuring the levels of ABA and GA during imbibition may help answer this question. Another non exclusive option would be that testa permeability to both exogenous germination inhibitors is increased in *tt15* seeds due to their flavanol defect. Indeed *tt4-8* mutant seeds deprived of flavonoids because of a mutation in the chalcone synthase gene (Supplemental Figure S2) exhibited the same dose-response curves to ABA and PAC than *tt15* alleles in Ws-4 (Figure 2D). Altogether these data suggest that *tt15* mutant seeds exhibit a germination syndrome with pleiotropic effects, which physiological analysis is complexified by altered seed coat permeability.

**Figure 2.**
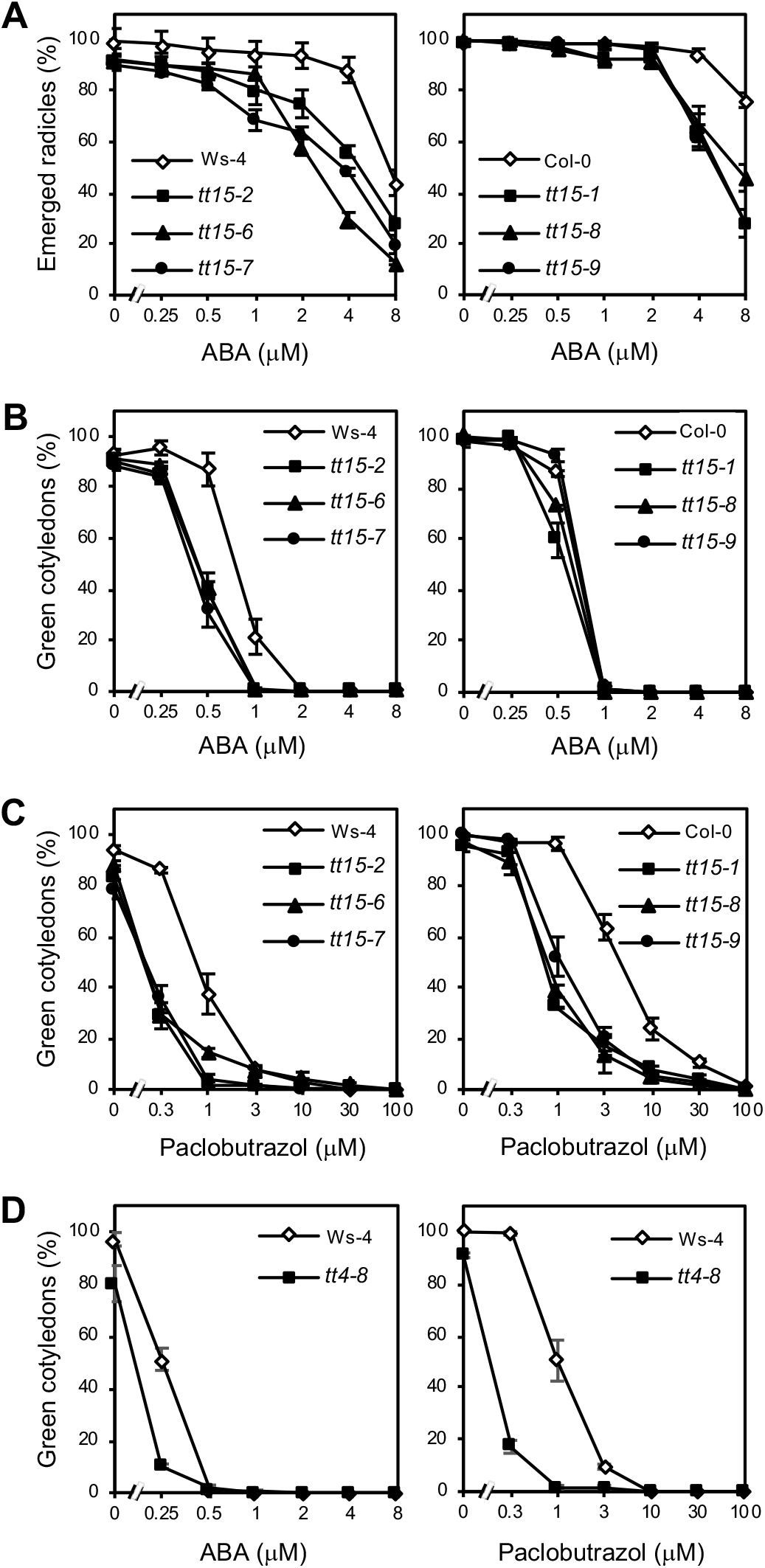
TT15 Depletion Increases Sensitivity of Seed Germination to Abscisic Acid and Paclobutrazol. **(A)** and **(B)** Sensitivity to exogenous abscisic acid (ABA), expressed as percentages of emerged radicles and green cotyledons, respectively. **(C)** Sensitivity to the gibberellin biosynthesis inhibitor paclobutrazol (PAC). **(D)** Sensitivity of *tt4-8* seeds to ABA and PAC. Error bars represent standard errors (n=3).

### Loss of TT15 Affects Endothelium and Endosperm Development

A histological analysis of developing seed structures was undertaken to understand further the mechanisms leading to the seed coat and seed lethality phenotypes in *tt15* mutants. Toluidine blue O (TBO) staining of class-I seed sections revealed that the *tt15-2* endothelium (ii1) layer had dramatically crushed and degenerated in the course of PC accumulation (Figure 3C compared to Figure 3A). PC stained dark blue with TBO fill the vacuole in wild-type endothelial cells (Figure 3B). However in *tt15-2* endothelial cells they appear as a flattened amorphous aggregate (Figure 3, D and F, arrowheads), suggesting that vacuole disruption may have occurred. Intermediate levels of ii1 cell degeneration, apparently taking place in a stochastic way, were observed (Figure 3, E and F, arrows). Aberrant endosperm cellularization and enlarged nodules especially at the chalazal pole were also observed in a few malformed (class III) *tt15-2* seeds around 6 days after flowering (late heart to torpedo stage of embryo development) when the endosperm is supposed to be cellularized (Figure 3, G and H). The same observation was made with *tt15-9* in Col-0 background (Supplemental Figure S8, A-D). A reduction of PC accumulation is not the cause of endothelium crushing because the endothelium of the chalcone synthase null mutant *tt4-8* (Ws-4 background) deprived of any flavonoid including PC exhibits well formed cells (Figure 3, I and J). Interestingly, the endothelium layer of the double mutants *tt15-2 tt4-8* (Figure 3, K and L) and *tt15-2 ban-1* accumulating anthocyanins in place of PC in endothelium (Supplemental Figure S9, C, D, G and H) is not crushed. Altogether these observations strongly suggest that flavanols (EC and/or PC) may specifically induce *tt15-2* endothelium cell death leading to precocious cell crushing. Whole-mount cleared seeds observed with differential interference contrast (DIC) microscopy revealed endothelial cell wall thickening and browning at the abaxial pole of the seed body (curving zone; Supplemental Figure S4A) in *tt15-2* seeds (Figure 3, M and N). Histochemical staining with the flavanol-specific reagent vanillin enabled to confirm the endothelial nature of the affected layer and confirmed cell wall thickening and browning both in *tt15-2* and *tt15-9* (Figure 3, O-Q; Supplemental Figure S8, E-G). As previously shown by Pourcel et al. (2005), flavanol oxidation in Arabidopsis seed coat forms brown products. It is therefore very likely that the brown reaction observed here is due to flavanol oxidation, especially as it is absent in *tt4-8* and *tt15-2 tt4-8* backgrounds (Supplemental Figure S9).

**Figure 3.**
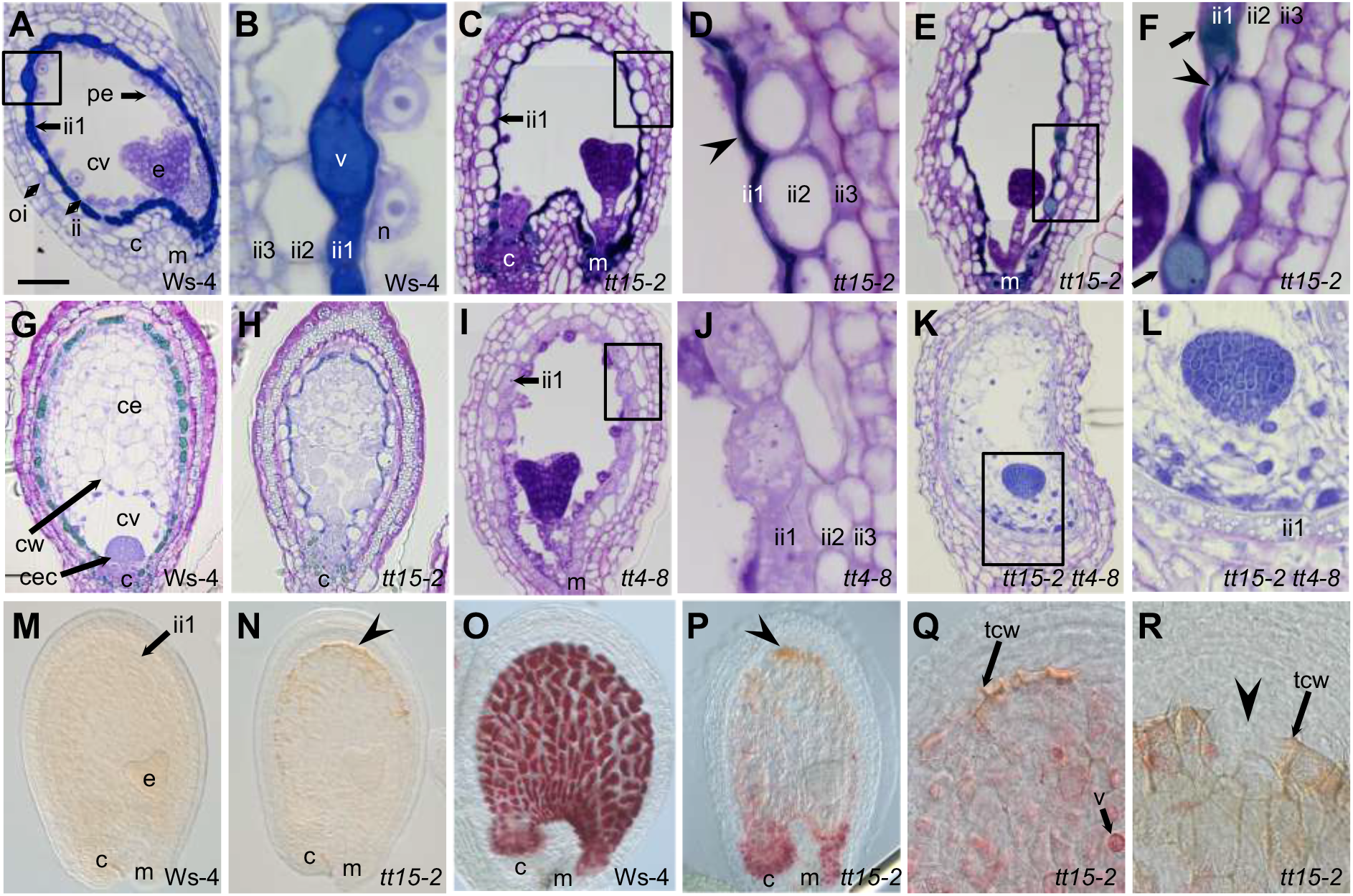
Loss of TT15 Affects Endothelium and Endosperm Development. **(A)** to **(L)** Longitudinal sections of developing seeds stained with toluidine blue O. **(A)** and **(B)** Dark blue-stained flavanols fill the wild-type vacuoles of endothelial cells (ii1 cell layer) at around four days after flowering (daf). **(C)** to **(F)** Most *tt15* endothelial cells flatten and degenerate in the course of flavanol accumulation (arrowheads). **(E)** and **(F)** A few ii1 cells exhibit unflattened cells with typical vacuolar structures (arrows) in some seeds. **(G)** Wild-type seed with a completely cellularized endosperm at around 6 daf. **(H)** Some *tt15* seeds (class III) display strong endosperm defects, with no obvious cellularization and an aberrant chalazal cyst. **(I)** and **(J)** Developing seeds of *tt4-8* deprived of flavonoids do not exhibit flatten endothelium. **(K)** and **(L)** In the double mutant *tt15-2 tt4-8* deprived of flavonoids, endothelium flattening and degeneration is suppressed compared with the situation in *tt5-2* background. **(B)**, **(D)**, **(F), (J)** and **(L)** are magnifications of insets from **(A)**, **(C)**, **(E), (I)** and **(K)**, respectively. **(M)** and **(N)** whole mount cleared seeds observed with differential interference contrast (DIC) microscopy. **(N**) Endothelial cells of *tt15* located at the abaxial pole of the seed (curving zone) exhibit oxidized (brown) PAs and cell wall thickening (arrowhead). **(O)** to **(R)** Detection of flavanols with vanillin staining (whole mounts at the heart stage; flavanols stain cherry red). **(O)** and **(P)** In *tt15* seeds, flavanols are present mainly at the micropyle and chalaza. Oxidized (brown) flavanols are observed at the abaxial pole (curving zone; arrowhead). **(Q)** and **(R)** Curving zone of developing *tt15* seeds at the heart stage stained with vanillin (whole mounts). Arrowheads show thickened brown endothelium cell walls **(Q)** and a breaking point in the endothelium layer **(R)**. C, chalaza; ce, cellularized endosperm; cec, chalazal endosperm cyst; cv, central vacuole; cw, cell wall; e, embryo; ii, inner integument; m, micropyle; oi, outer integument; n, nodule; tcw, tannic cell wall; v, vacuole. Bar in **(A)** = 50 μm in **(A)**, **(C)**, **(E)**, **(G)**, **(H)**, **(I)**, **(K)** and **(M)** to **(P),** 10 μm in **(B)**, **(D)**, **(F)**, **(J)** and **(L),** and 25 μm in **(Q)** and **(R)**.

Some seeds exhibit endothelium rupture probably due to exacerbated mechanical tension after cell wall overthickening (Figure 3R; Supplemental Figure S8H). This polarized thickening at the abaxial side of the seed body defines a weakness zone which may enable the expanding embryo to be expelled from the testa (Figure 1D). To analyze further the cellular mechanisms at play during endothelium polarized modification, we performed a histochemical detection of reactive oxygen species (ROS) in developing seeds using the fluorochrome DCFH-DA. Interestingly ROS were present and shown to accumulate essentially at the abaxial pole of the endothelium (Figure 4, A and B). During immune response, ROS such as H_2_O_2_ are known to trigger the accumulation of callose, a β-1,3-linked glucose polymer (Luna et al., 2011). Here, using aniline blue staining, we could also detect ectopic callose deposition in endothelium (Figure 4, C and D).

**Figure 4.**
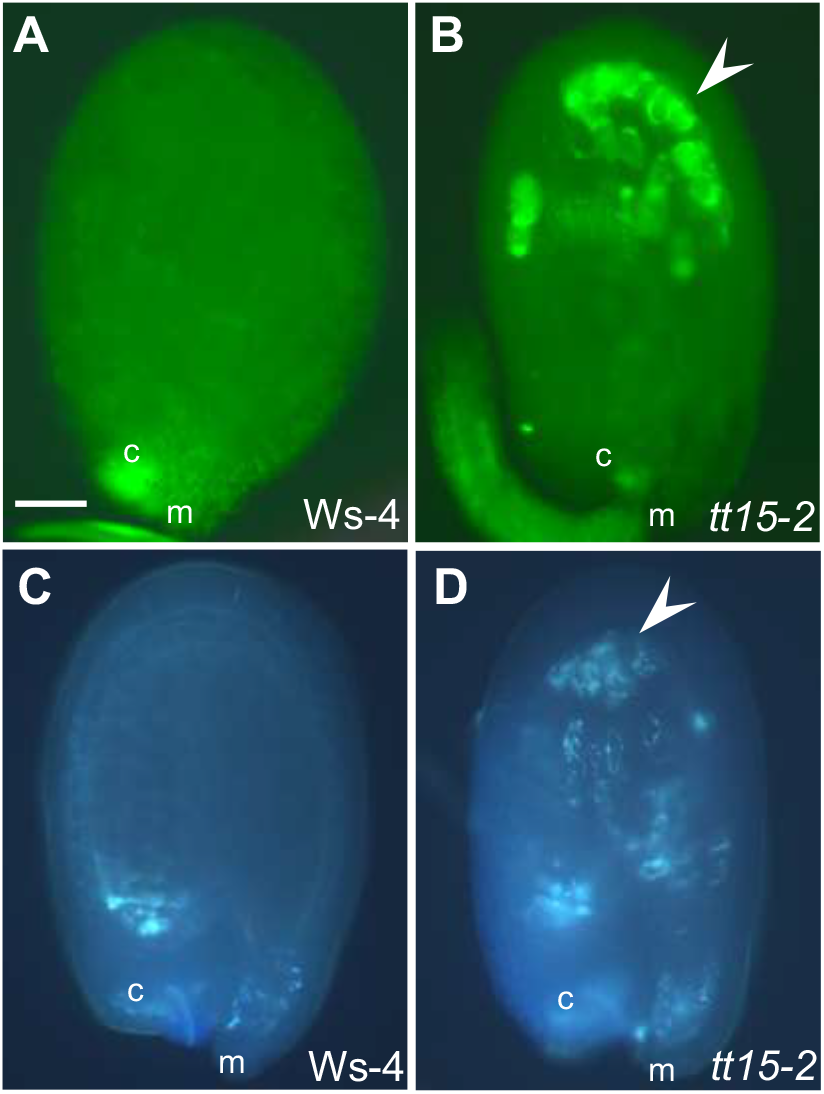
*TT15* Disruption Triggers an Autoimmune-like Response in Seed Coat Endothelium. Developing seeds observed under UV light are shown (whole mounts). **(A)** and **(B)** Detection of ROS with DCFH-DA staining. Arrowhead shows ROS accumulation polarized at the seed abaxial side. **(C)** and **(D)** Detection of callose with aniline blue staining. Arrowhead shows ectopic callose deposits. c, chalaza; m, micropyle. Bar in **(A)** = 50 μm in **(A)** to **(D).**

### Spatio-Temporal Analysis of Promoter Activity and mRNA Shows that TT15 is Expressed in Reproductive and Vegetative Tissues

A 2.0-kb DNA sequence upstream of the ATG translation codon was used to establish the spatio-temporal pattern of *TT15* promoter activity in Arabidopsis Ws-4 stable transformants expressing the *pTT15:uidA* construct (Figure 5). As expected from *tt15* mutant seed phenotypes, promoter activity was detected in endothelium (Figure 5, A, J and K), embryo (Figure 5, B, C and L), endosperm (Figure 5, I and L), germinating seeds and young seedlings (Figure 5, F, G and H). Activity was also detected in mucilage layer (oi2 outer integumentary layer; Figure 5, B, J and K), chalaza-funiculus continuum (Figure 5, A and J), seed abscission zone (Figure 5, B-D) and unfertilized necrotic ovules (Figure 5E). As shown in Supplemental Figure S10, the *TT15* promoter was activated in other reproductive organs (ovule primordia and ovules, gynoecium, style, transmitting tract, nectaries and pollen grains) and also in various vegetative tissues (cauline and root meristems, lateral root primordia, root cap, vascular bundles from cotyledons, leaves, roots and stems, stomata, and hydathodes). The stronger *tt15* mutant phenotypes observed in Ws-4 background compared with Col-0 prompted us to perform a quantitative polymerase chain reaction (qPCR) analysis of *TT15* expression in both accessions (Supplemental Figure S11). This analysis did not reveal any significant differences except in the amount of stored mRNA in dry seeds, which was higher in Ws-4 than in Col-0. A peak of expression was also observed in senescing siliques of both accessions. Interestingly *TT15* mRNA stored in mature seed disappears during imbibition until resuming when early seedling growth occurs. Altogether, our experimental data corroborated by public transcriptomes (Supplemental Figure S12) strongly argue towards a role for TT15 not only in seed development and germination but also in seedling growth.

**Figure 5.**
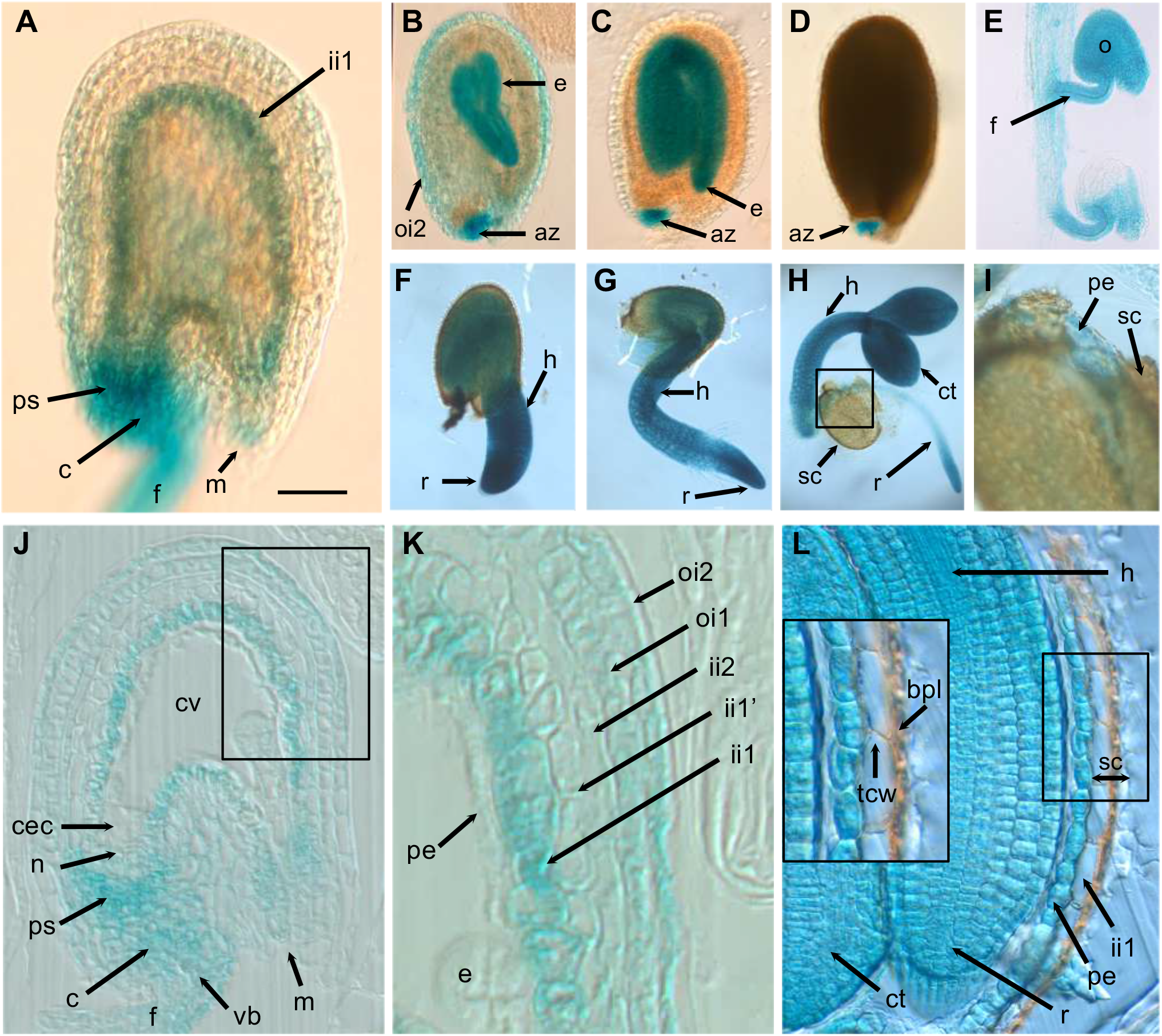
The *TT15* Promoter is Active during Seed Development and Germination. Arabidopsis transformants expressing the *ProTT15:uidA* construct were analyzed for GUS reporter activity. A 2.0-kb sequence upstream of ATG was used as promoter. **(A)** to **(D)** Developing seeds at the globular **(A)**, torpedo **(B),** cotyledonary **(C)** and late maturation **(D)** stages (whole mounts). **(E)** Unfertilized ovules (aborted seeds) from a developed silique (whole mounts). **(F)** to **(I)** Germinating seeds at 32 h **(F),** 48 h **(G)** and 72 h **(H)** after imbibition; **(I)** Magnification of the box in **(H)**. **(J)** Section of developing seed at the quadrant stage. **(K)** Magnification of the box in **(J)**. **(L)** Longitudinal section of a mature seed. The inset is a magnification of the boxed area. az, abscission zone; bpl, brown pigment layer; c, chalaza; cec, chalazal endosperm cyste; ct, cotyledon; cv, central vacuole; e, embryo; f, funiculus; h, hypocotyle; ii, inner integument; m, micropyle; n, nucellus; o, ovule; oi, outer integument; pe, peripheral endosperm; ps, pigment strand; r, radicle; sc, seed coat; tcw, tannic cell wall; vb, vascular bundle. Bar in **(A)** = 50 μm in **(A)** and **(J),** 140 μm in **(B)** to **(D),** 80 μm in **(E),** 250 μm in **(F),** 350 μm in **(G),** 550 μm in **(H),** 200 μm in **(I),** 25 μm in **(K)**, and 50 μm in **(L) (**20 μm in inset).

### TT15 Co-localizes with Tonoplast and Amyloplast Markers and is also Present in Cytoplasm

To determine the subcellular localization of the TT15 protein, transgenic Arabidopsis plants expressing the green fluorescent protein (GFP)-TT15 and TT15-GFP constructs placed under the control of a dual 35S promoter were generated (Supplemental Figure S13A). Both constructs complemented *tt15-2* seed coat phenotypes, *i.e.* rescued wild-type seed coat colour (Supplemental Figure S13B) and vacuolar deposition of flavanols was restored in the transformants (Supplemental Figure S13C). Seed flavonoid profiling of the *tt15-2* allele showed a strong reduction of flavanols, including epicatechin hexoside (EC-H; Supplemental Figure S13D), which is consistent with the results from the vanillin histological assay. A significant reduction in the flavonol quercetin 3-O-rhamnoside or QR (Supplemental Figure S13D) was also observed. We detected minor variations in other flavonols as well. All modifications were rescued in transformants. Supplemental Table S2 displays the complete list of flavonoid compounds identified by our UPLC-MS analysis in wild-type mature seeds. After having checked that the same localization pattern was observed with TT15-GFP (Supplemental Figure S14), the following part of our study focused on GFP-TT15.

Confocal imaging microscopy of GFP-TT15 in developing seeds was realized on the mucilage layer (oi2) of developing seed coat. Imaging at the level of the endothelium (ii1) was not possible due to a weak activity of the 35S promoter in this cell layer. The presence of GFP fluorescence was revealed at the tonoplast, at localized regions of amyloplasts and in the cytosol (Figure 6, A-C; Supplemental Figure S14A). In cotyledon epidermis, tonoplast fluorescence was particularly conspicuous at the level of stomata, probably because of the presence of numerous vacuolar convolutions in guard cells (Figure 6D; Supplemental Movie S1). However no obvious signal was observed at the level of chloroplasts. GFP-TT15 at the tonoplast was also observed in seedling roots (Supplemental Figure S14, E-G). As imaged in Supplemental Movies S2 and S3, GFP-TT15 was also located in mobile aggegates or vesicles of various sizes present in the cytosol and also in vacuolar lumen where their movement seemed to be constrained by the tonoplast. The identity of these structures remains to be elucidated.

**Figure 6.**
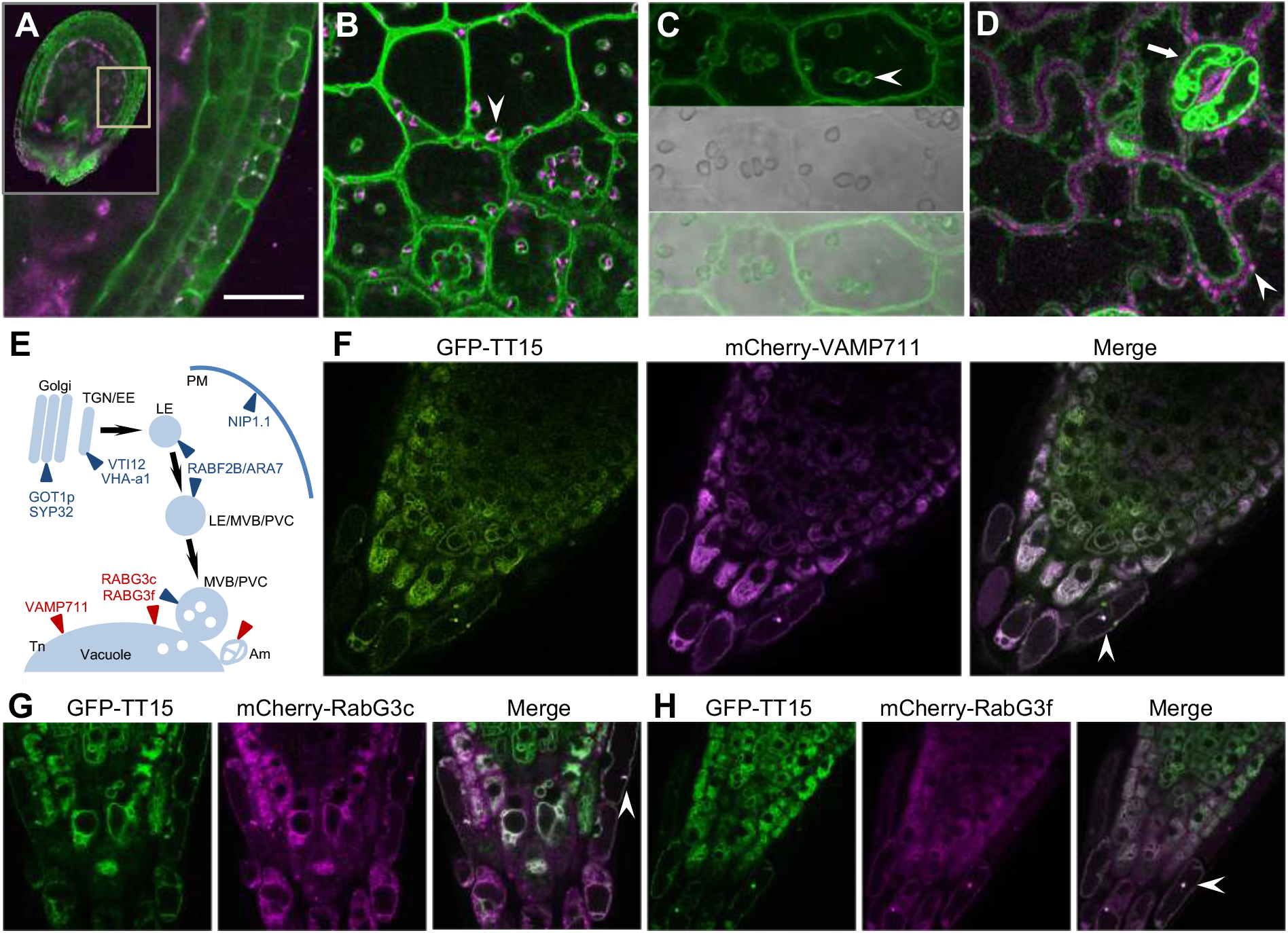
GFP-TT15 Localizes at Tonoplast, Amyloplasts, Cytoplasm and Unknown Intravacuolar Aggregates. Confocal fluorescence micrographs of Arabidopsis stable transformants are shown. **(A)** Transverse optical section of a developing seed at the globular stage expressing GFP-TT15, with magnification at the level of integuments (inset). **(B)** and **(C)** Top view of a seed coat epidermis (oi2 cell layer) at the early heart stage of embryo development (around 4 days after flowering) expressing GFP-TT15. Arrowheads show amyloplasts partially co-localizing with GFP-TT15. **(C)** is part of Supplemental Figure 16A (inset). **(C)** Cotyledon epidermis from a 3-day-old seedling expressing GFP-TT15, with a stomata (arrow) and chloroplasts (arrowhead). Magenta in **(A)**, **(B)** and **(D)** reveals autofluorescence. **(D)** Scheme of subcellular trafficking pathways showing the location of markers used in this study. In red and blue are markers that co-localize or do not co-localize with GFP-TT15, respectively. **(E)** to **(H)** Co-localization of GFP-TT15 with various subcellular compartment markers on 5-day-old roots are shown. Arrowheads point to cells with co-localizing intravacuolar aggregates. **(F)** Tonoplast (Tn); **(G)** and **(H)** Multivesicular bodies (MVB) / prevacuolar compartments (PVC). Bars = 10 μm. Am, amyloplast; Tn, tonoplast.

To progress further in the identification of GFP-TT15 intracellular locations, fluorescent markers for various subcellular compartments were introduced in GFP-TT15 transformant background by crosses (Figure 6E; Supplemental Table S4). GFP-TT15 fluorescence co-localized with the tonoplastic marker mCherry-VAMP711 in epidermis, columella and lateral root cap of seedling roots (Figure 6F). It also partially co-localized with markers mCherry-RabG3c and mCherry-RabG3f at the level of tonoplast but not at the level of multivesicular body (MVB)/prevacuolar compartments (PVC; Figure 6, G and H). On the other hand, no co-localization of GFP-TT15 was observed with Golgi apparatus, trans Golgi netwok (TGN)/early endosome (EE), late endosome (LE)/multivesicular body (MVB)/prevacuolar compartment (PVC) and plasma membrane (PM) (Supplemental Figure S15). Intriguingly, the intravacuolar mobile aggregates identified by GFP-TT15 in roots, as mentioned above, happened to co-localize with all our subcellular compartment markers (Figure 6, F-H, white arrowheads; Supplemental Figure S14, A-F, white arrowheads).

### Alteration of Vacuole Development in Endothelium Causes a *transparent testa* Phenotype

We have shown that *tt15-2* seeds at the heart stage of embryo development (Figure 3P) exhibit endothelial cell breakdown causing a strongly reduced flavanol accumulation, compared with wild-type endothelial cells having central vacuoles filled with flavanols (Figure 3O). We failed to detect any significant defect in expression of flavonoid biosynthetic and regulatory genes in developing seeds (Supplemental Figure S16), which suggests that the lower amount of accumulated flavanols in *tt15-2* endothelial cells is rather due to abnormal cell development. To progress further in the identification of the disrupted cellular functions in *tt15-2*, we observed the dynamics of vacuole morphogenesis in endothelial cells during seed development using vanillin-stained flavanols as vacuolar markers (Figure 7). The early spatio-temporal pattern of flavanol deposition is similar between *tt15*-2 and wild type (Figure 7, A and E). A difference becomes visible at the globular stage, with endothelial cells stopping vacuole filling and starting a progressive breakdown (Figure 7, B-D and F-H). We named this degeneration process PECD for Premature Endothelium Cell Death, to distinguish it from Programmed Cell Death (PCD) occurring in wild-type endothelium at later stages of seed development, as described by Andème Ondzighi et al. (2008). The PECD syndrome is visible first, and is the most intense at the abaxial pole of the seed (curving zone). From the heart stage onwards, most *tt15-2* endothelial cells (region 2; Supplemental Figure S4A) look empty (Figure 7H) by comparison with wild-type cells filled with flavanols (Figure 7D). Only the micropyle (region 1) and chalaza (region 3) exhibit flavanols in *tt15-2* seeds. We observed frequently a few endothelial cells that, apparently with a stochastic pattern, exhibited transient flavanol accumulation. Subcellular organization at the octant stage is similar between both genotypes (Figure 7, I and M). Vacuolar morphology is predominantly roundish with intensely stained tonoplast and intravacuolar aggregates. Differences beween *tt15-2* and wild type are obvious from the globular stage onwards (Figure 7, J-L and N-P). Wild-type vacuole morphology evolves from roundish to elongated. Vacuole lumen accumulates substructures resembling small vesicles with stained membranes that start to accumulate flavanols (Figure 7, J and K). At the heart stage, these substructures fill completely the endothelial vacuole, which volume is constrained by the surrounding cell wall (Figure 7L). In *tt15-2* mutant, vacuole morphogenesis is blocked at the globular stage (Figure 7N). Indeed the roundish vacuoles do not elongate and very limited substructural organization of the lumen is taking place. Different stages of evolution can be observed in the same endothelium. The degeneration process ends with cell death as suggested by vacuole disappearance in most endothelial cells at the heart stage.

**Figure 7.**
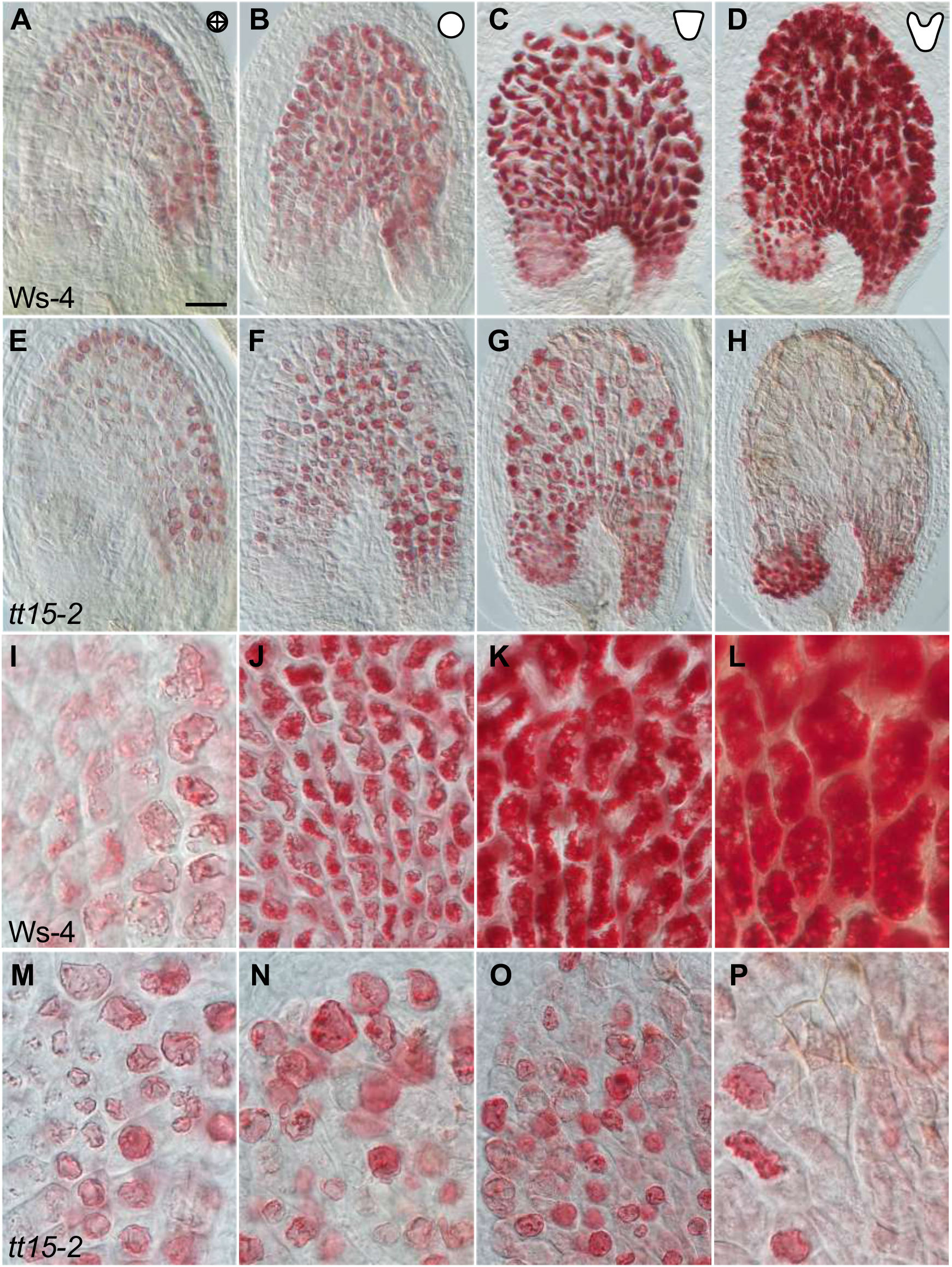
Alteration of Vacuole Development in Endothelium Causes a *transparent testa* Phenotype. The spatio-temporal pattern of vacuole dynamics in endothelial cells is monitored by following flavanol deposition in vacuoles. Colorless flavanols stain cherry red with vanillin (whole mounts). **(A)** to **(H)** Premature endothelium cell death (PECD) is observed in *tt15-2* mutant seeds from around the globular stage **(F)** onwards. In comparison, wild-type endothelial vacuoles are filled with flavanols at the heart stage **(D)**. **(I)** to **(P)** Vacuole shape and lumen organization are modified in developing endothelial cells of *tt15-2* mutant seeds. Bar in **(A)** = 40 μm in **(A)**, **(B)**, **(E)** and **(F)**, 50 μm in **(C)**, **(D)**, **(G)** and **(H)**, 6 μm in **(I)**, **(L)**, **(M), (N)** and **(P)**, 7 μm in **(J)**, **(K)**, and **(O)**.

### Tonoplast Fluidity is Increased in *tt15* Roots

Knowing that SG and ASG are components of plant tonoplasts (Yoshida and Uemura, 1986; Tavernier et al., 1993; Yamaguchi and Kasamo, 2001) and that they have the ability to efficiently order membranes and thus modulate their fluidity or viscosity (Laloi et al., 2007; Halling et al., 2008; Grosjean et al., 2015) prompted us to investigate whether *tt15*-2 tonoplast fluidity was modified. Fluorescence recovery after photobleaching (FRAP) was carried out to quantitatively monitor the lateral diffusion characteristics of the tonoplast in epidermis of Arabidopsis seedling roots (Figure 8). Membrane labeling was done with the fluorescent probe GFP-NRT2.7 (Supplemental Table S4). NRT2.7 is an Arabidopsis nitrate transporter located specifically at the tonoplast (Chopin et al., 2007). The probe was introduced in *tt15-2* background by crossing. After having checked that no alteration of GFP-NRT2.7 location was observed in the mutant compared with the wild type (Figure 8A), photobleaching was performed with both genotypes at the tonoplast of cells from the root elongation zone (Figure 8B). Two parameters t_1/2_ (recovery half-life) and M_f_ (mobile fraction) were used to describe membrane lateral mobility in quantitative analysis (Figure 8, C and D). The recovery half-life t_1/2_ provides a measure of the half-life recovery time and M_f_ indicates the fraction of fluorescent molecules recovered into the bleached region. As shown in Figures 8C and 8D, the level of GFP-NRT2.7 was restored to 83% at 40 s in the bleached ROI in *tt15-2* mutant (M_f_=83%; Linf=67% and Lsup=96%), while there was a 79% recovery in that of wild type Ws-4 (M_f_=79%; Linf=63% and Lsup=95%). These results revealed that a significantly lower proportion (17%) of GFP-NRT2.7 was present as the immobile fraction (I_f_) in *tt15-2* mutant as compared with the 21% in wild type. A significant difference was also observed for the recovery rate, with t_1/2_ of mutant and wild type being 4.505 s and 6.875 s, respectively. Altogether these data demonstrate that decreasing the level of SG increases tonoplast fluidity, thus alters its dynamics.

**Figure 8.**
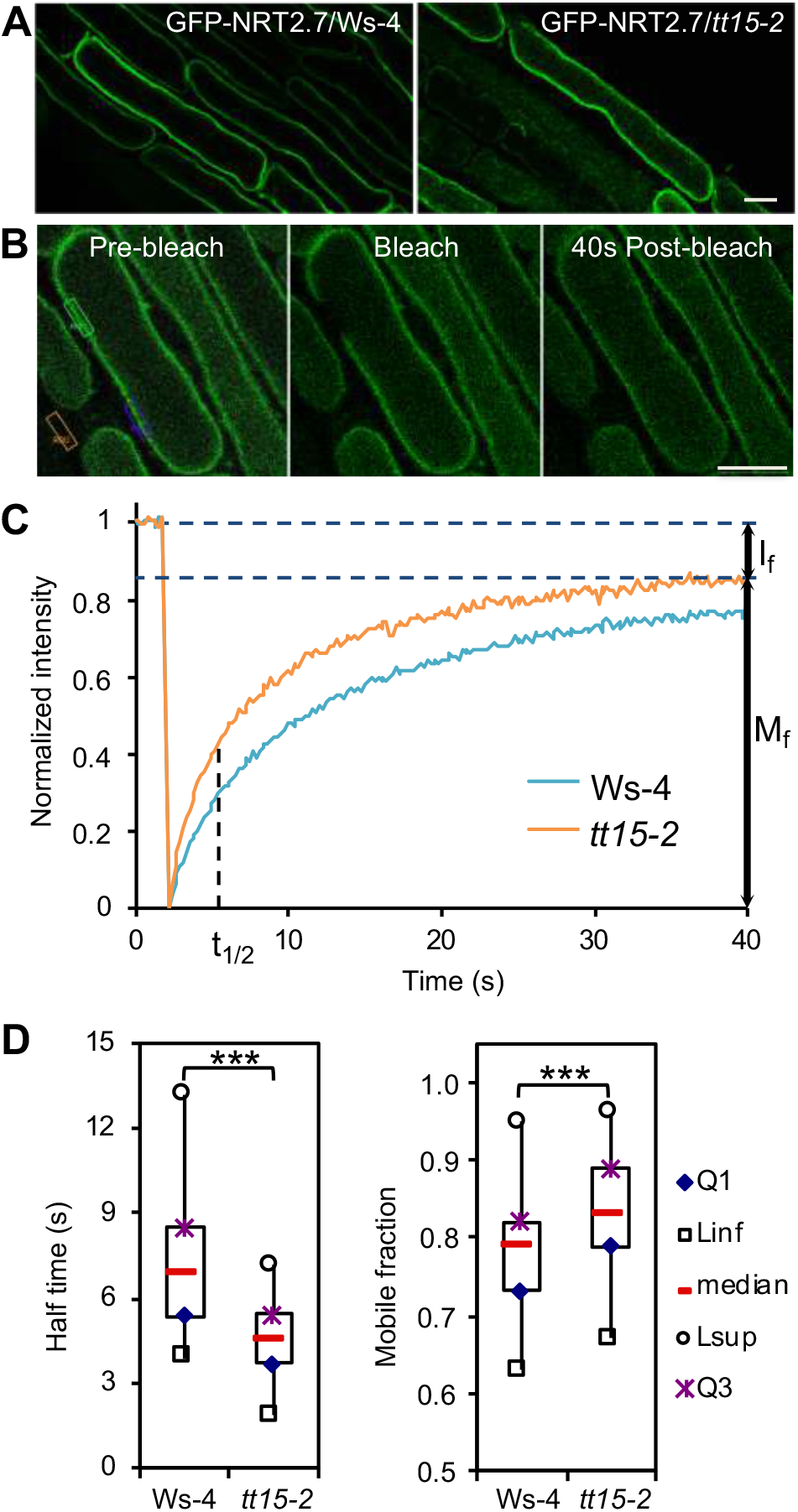
Tonoplast Membrane Fluidity is Increased in *tt15* Mutant Background. Fluorescence recovery after photobleaching (FRAP) of GFP-NRT2.7 was performed at elongating root epidermal cells of 4 day-old seedlings from Ws-4 wild type and *tt15-2* mutant. **(A)** GFP-NRT2.7 subcellular localization is not modified in *tt15-2* background. **(B)** Fluorescence-intensity imaging prior to, immediately following, and 40 s after photobleaching. Region of interest 1 (ROI1): bleached area (green); ROI2: unbleached area (purple); ROI3: background area (orange). **(C)** Quantitative FRAP analysis. Median values of normalized fluorescence intensities of 66 measurements for each genotype are shown. M_f_, mobile fraction; I_f_, immobile fraction; t_1/2_, recovery half-time. **(D)** Box plots of median values of normalized and fitted data for recovery half time and mobile fraction are shown. Asterisks indicate significant differences (P < 0.001) between samples by Wilcoxon test. Bars = 10 μm in **(A)** and **(B)**.

### The *tt15* Mutation Partially Rescues the Vacuolar Phenotype of *tt9* in Endothelium

Looking for functional relationships between TT15 and other TT proteins which mutants have a similar pattern of flavanol deposition in the seed coat, namely TT9, TT1 and TT16, may put some light on the mechanisms involved in *tt15* endothelium. Here, we focused on TT9 because, as a peripheral membrane protein involved in vacuolar development and trafficking (Ichino et al., 2014), it appeared the most likely candidate to fulfill our objectives.

As shown above, *tt15-2* (Figure 1B) and *tt9-1/gfs9-4* (Supplemental Figure S4C) exhibit a similar patterning of seed coat pigmentation, with only the endothelium being defective in PC accumulation. Interestingly whole-mount seed clearing (Figure 9B), TBO-stained sections of developing seeds (Figure 9D) and vanillin assay for flavanol detection (Figure 9E) also revealed strong similarities between *tt9-1* and the *tt15-2* phenotypes presented in Figure 3, notably endothelium degeneration in the course of flavanol accumulation ending with endothelium cell death together with cell wall thickening and browning at the curving zone. (Ichino et al., 2014) previously revealed that *tt9* mutants exhibit vacuole fragmentation using light microscopy and transmission electron microscopy analysis of endothelium. Here, this vacuolar phenotype could be confirmed through the histological detection of flavanols using TBO (Figure 9D arrow) and vanillin (Figure 9, E and G). We did not observe vacuolar fragmentation with *tt15* mutants (Figure 3H and Supplemental Figure S17). These observations prompted us to determine the epistasis relationships between both mutations by constructing the double mutant *tt15-2 tt9-1*. Double mutant seeds exhibited a novel seed coat pigmentation (Figure 9L compared with Figure 9, I-K) with an absence of differential patterning between the micropyle-chalaza region and the seed body and a different overall colour. Importantly the *tt9-1* mutation suppressed the *tt15-2* PECD phenotype (Figure 9F). Moreover a novel vacuolar morphology and luminal organization could be observed (Figure 9H), where flavanols are detected only in vesicle-like structures aggregating outside the tonoplast of medium-sized vacuoles. Altogether these data demonstrate that TT15 and TT9 genetically interact in the sense that they both contribute to vacuole development, however in partially overlaping biochemical pathways.

**Figure 9.**
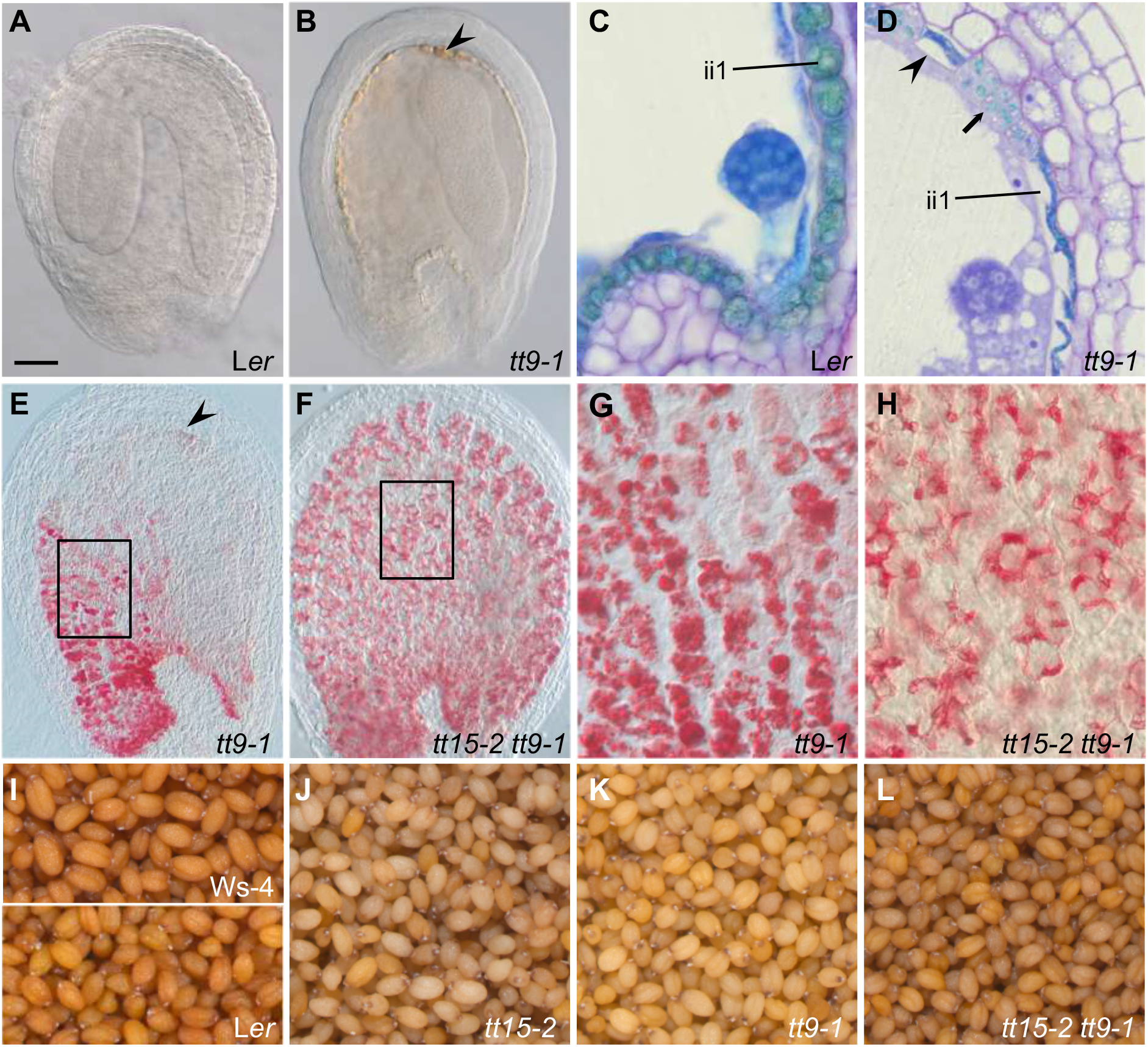
TT15 and TT9/GFS9 Genetically Interact. **(A)** to **(D)** The *tt9-1/gfs9-4* mutant endothelium phenotype resembles *tt15-2* phenotype. **(A)** Developing wild-type L*er* seed cleared in chloralhydrate solution (whole mount) **(B)** Developing seed of *tt9-1* mutant cleared in chloralhydrate solution (whole mount). Endothelial cells exhibit oxidized (brown) PCs and cell wall thickening (arrowhead) mainly at the curving zone (arrows). **(C)** and **(D)** Longitudinal sections of developing seeds stained with toluidine blue revealing flavanols in greenish blue. **(D)** As for *tt15-2,* most *tt9-1* endothelial vacuoles flatten and degenerate in the course of flavanol accumulation (arrowhead). Some cells show fragmented vacuoles typical for *tt9-1* (arrow). ii1, inner integument 1 (endothelium). **(E)** to **(H)** The *tt9-1* mutation partially rescues the *tt15-2* endothelium defects. Detection of flavanols was done with vanillin staining in developing seeds at the heart stage (whole mounts). **(E)** In *tt9-1* seeds as for *tt15-2*, PCs are present mainly at the micropyle and chalaza and oxidized (brown) PCs are observed at the abaxial pole (curving zone; arrowhead), due to precocious endothelial cell death (PECD)**. (F)** PECD is not observed in the double mutant. **(G)** and **(H)** are magnifications of endothelial cells showing vacuole and vesicle morphology (insets in **(E)** and **(F)** point to the respective locations). **(I)** to **(L)** Mature seed colours. The double mutant *tt15-2 tt9-1 e*xhibits a novel pigmentation phenotype. Bar in **(A)** = 50 μm in **(A), (E)** and **(F)**, 25 μm in **(B)** to **(D)**, 6 μm in **(G)**, 7 μm in **(H)**, and 500 μm in **(I)** to **(L).**

## DISCUSSION

Here we have identified a function for the sterol-3-β-glucosyltransferase TT15 in regulating vacuolar membrane characteristics and vacuole functions in Arabidopsis. In addition to revealing a seed lethality phenotype with variable penetrance and an increased sensitivity of seed germination to exogenous inhibitors, our work brought novel informations on the cellular mechanism causing the pale seed coat colour phenotype and revealed a genetic interaction between TT15 and TT9, a protein involved in vacuolar biogenesis and vesicular trafficking. The analysis of an allelic series of six *tt15* mutants in two different accessions was important to confirm the robustness of the observed phenotypes and to determine which part of the variations was imparted to the genetic background. Taken together, our data suggest an involvement of the vacuole, SG and flavonoids in the observed traits, as discussed below.

### TT15 is Essential for Seed Development and Germination

A seed lethality phenotype exhibiting incomplete penetrance and maternal inheritance was observed for both *tt15-2* and *tt15-9* mutant alleles in Ws-4 and Col-0 backgrounds, respectively. Moreover an endosperm cellularization defect was detected in the most affected seeds. The percentage of aborted seeds was significantly higher in Col-0 than in Ws-4 background. Moreover the proportion of dead seeds was non-mendelian, modulated by environmental conditions and higher when the *tt15-2* mutation was transmitted by the female parent. This developmental syndrome reminds lethality of Arabidopsis F1 hybrids caused by interploidy and interspecific crosses between diverged parents, namely the triploid block. The endosperm plays a central role in this gene dosage-sensitive incompatibility, together with the maternally expressed WRKY transcription factor TTG2 which disruption suppresses the triploid block, decreases endothelial cell elongation leading to smaller seeds and causes precocious endosperm cellularization (Garcia et al., 2005; Dilkes et al., 2008; Burkart-Waco et al., 2013; Köhler et al., 2021). The *tt4* and *tt8* mutations affecting chalcone synthase and a bHLH transcription factor, respectively, also act as maternal suppressors of triploid block (Buer and Muday, 2004; Doughty et al., 2014; Zumajo-Cardona et al., 2023), which points to endothelial flavanols as potential inducers of seed lethality in interspecies and interploidy crosses. As *tt15* mutants have small seeds (DeBolt et al., 2009)(this study), a situation mimicking to some extent a maternal-excess scenario where maternally-expressed genes are less imprinted and thus are more expressed in the endosperm is likely and would be consistent with the fact that the mutation is less transmitted through the ovule. However endosperm with defective cellularization reminds a paternal-excess situation, which suggests that genes overexpressed on the paternal side may also interfer with seed development in absence of TT15. The fact that *TT15* is expressed both in the endosperm and the seed coat (this study) complexifies the understanding of the *tt15* seed development phenotypes.

The moment at which the endosperm cellularizes after a phase of free nuclear divisions without cytokinesis (endosperm proliferation) is a crucial determinant of seed size (Sorensen et al., 2002; Hehenberger et al., 2012; Doughty et al., 2014). Upon fertilization, the auxin phytohormone produced in the central cell of the embryo sac and afterwards in endosperm triggers seed coat differentiation (Figueiredo et al., 2016). Increased biosynthesis and signalling of auxin in the endosperm prevents its cellularization and leads to seed developmental arrest, thus phenocopying the phenotype of paternal-excess triploid seeds (Batista et al., 2019). Moreover flavonoids are negative regulators of auxin transport (Buer and Muday, 2004). In this context, we hypothesize that small seed size, seed lethality and endosperm cellularization defects observed in *tt15*-2 may be correlated with a perturbation of auxin homeostasis. Relevant with this hypothesis is the presence of seedlings with abnormal cotyledon number in the progeny of some *tt15-2* plants revealing stem cell niche perturbation (this study). Such a phenotype is regularly observed in situations when auxin intracellular transport is affected, as for instance in mutants affected in sterol homeostasis (Souter et al., 2002), in vacuolar sorting of auxin carriers (Jaillais et al., 2007) or in auxin-mediated ribosomal biogenesis regulating vacuolar trafficking (Rosado et al., 2010). It would be interesting to investigate by a microscopic approach whether auxin sensors such as *DR5-uidA* are deregulated in *tt15* developing seeds.

Seeds of *tt15* exhibit a pleiotropic germination syndrome. Consistent with the situation prevailing with mutants which seed coat is affected in flavanol metabolism (Debeaujon et al., 2000) and/or lipid polyester (cutin, suberin) deposition (Molina et al., 2008), previous studie have shown that *tt15* mutant seeds exhibit a reduced seed dormancy and an increased testa permeability to tetrazolium salts (Focks et al., 1999; DeBolt et al., 2009; MacGregor et al., 2015; Loubéry et al., 2018). In our laboratory conditions and with all our alleles in two accession backgrounds (Col-0 and Ws-4) we could confirm a reduced primary dormancy and an increased testa permeability to tetrazolium salts. Moreover we observed an hypersensitivity to the germination inhibitors ABA and PAC brought exogeneously, as previously reported for several other *tt*s (Debeaujon and Koornneef, 2000; Tamura et al., 2006). Increased testa permeability to ABA and PAC may be responsible for this phenotype, especially knowing that *tt15* cumulates defects in both flavanol and lipid polyester metabolisms (DeBolt et al., 2009). Already we showed here that ABA levels in dry seeds are unmodified. Our analysis also showed the existence of a natural variation for ABA content in dry seeds, with Ws-4 having more ABA than Col-0.

### TT15 disruption Causes Premature Endothelium Cell Death

Wild-type Arabidopsis endothelium undertakes PCD, starting at the torpedo embryo stage and being effective around the bent-cotyledon stage. It progresses from the abaxial zone (curving zone) towards chalaza and micropyle, as the cellular endosperm expands (Andème Ondzighi et al., 2008). Here, we revealed that the *tt15* endothelium exhibits PECD, a precocious degeneration being visible from the globular stage onwards and ending in cell death. Contrarily to the situation observed in WT PCD, the *tt15* endothelium layer completely collapses, with the exception of a few cells due to a weak penetrance of the phenotype. The cellular mechanisms associated with ii1 premature cell death progression from the globular stage of embryo development onwards involve the interruption of vacuole development followed by vacuolar collapse probably due to tonoplast lysis. The vacuole is a central player in the execution of cell death (Shimada et al., 2018). Under this scenario, flavanols that have started to accumulate in ii1 vacuoles may leach into the cytosol and migrate towards the cell walls where they would be oxidized into brown products possibly by resident laccases and peroxidases and cross-link with polysaccharides (Pourcel et al., 2005).

We observed premature degeneration leading to cell death at the abaxial pole of the *tt15* endothelium, which was correlated with H_2_O_2_ production and callose deposition. This cellular response reminds the signature of autoimmunity or lesion mimic syndrome triggered by a spontaneous deregulation of nucleotide-binding domain leucine-rich repeat (NLR) receptors that are normally engaged in effector-triggered immunity upon pathogen attack (Ben Khaled et al., 2015; Freh et al., 2022). The trigger of this polarization may be mechanical stress at the level of the curving zone to which the *tt15* mutant endothelium would respond by exacerbated cell death and wall thickening. Creff et al. (2015) showed that the adaxial epidermis of the outer integument (oi1 cell layer) is a mechanosensitive cell layer responding to the mechanical stress exerted by the expanding embryo and endosperm by cell wall thickening. In this context, we speculate that the endothelium or adaxial epidermis of the inner integument (ii1 cell layer) may also be a mechanosensitive cell layer responding to filial tissue expansion, which pressure would be higher at the curving zone. The reason why this response is exacerbated and leads to a HR-type of cellular mechanism in a *tt15* background deserves to be explored further. On the same line, Burkart-Waco et al. (2013) reported that a perturbation of the communication between endosperm and maternal tissues in Arabidopsis interspecific crosses caused the activation of defense-like responses. Perturbation of *tt15* tonoplast homeostasis through its modified SG composition may possibly affect its resistance and/or the function of resident proteins such as the tonoplast-located mechanosensor PIEZO (Radin et al., 2021). As a corollary, 1-2% *tt15* mature seeds exhibits some vivipary, with an atypical emergence of the embryo from the seed coat at the abaxial pole of the seed suggesting perturbed cell wall integrity at the curving zone. We hypothesize that due to increased thickening caused by callose deposition, the *tt15* cell wall looses its extensibility and thus its resistance to embryo growth, creating a weakness zone.

Previous works demonstrated that recombinant TT15 could catalyze the glucosylation of free sterols *in vitro*, which was relevant with a reduction of glucosylated sterols in *tt15* mature seeds (DeBolt et al., 2009; Stucky et al., 2015) (this study). Perturbation of sterol homeostasis is therefore likely to explain the pleiotropic phenotypes of the *tt15* mutants. Sterols serve multiple biological functions, from structural components of membranes to signaling molecules as precursors of BR (Clouse, 2002; Zhang et al., 2015). Shimada et al. (2021) observed that excess sterol led to the development of a darker seed coat due to proanthocyanidin overaccumulation. Cell death-mediated shrinkage of the inner integument was also impaired, resulting in a thicker seed coat and a delayed seed germination. Intriguingly, these phenotypes seem opposite to the ones observed with the *tt15* mutant, which may be explained by the fact that excess free sterols that are toxic to the cell machinery may possibly be neutralized through glucosylation by TT15 (Supplemental Figure S1). Altogether, these observations suggest the existence of a complex interplay between the developing endothelium and endosperm involving TT15, SG and flavanols, which perturbation causes endothelium degenerescence and endosperm cellularization defects in a gene- or presumably auxin signal-dosage-dependent manner. In this context, it will be worse investigating whether TT15 works as a hub to integrate auxin-regulated developmental program and vacuole trafficking through the regulation of sterol metabolism, similarly to the situation observed for the ribosomal protein RPL4 by Li et al. (2015).

### TT15 is Mainly Located at the Tonoplast

Knowing the precise subcellular localization of SGTs is crucial for a thorough understanding of their biological functions. Previous studies in various plant species and with diverse experimental approaches reported multiple subcellular localizations for SGTs, including cytoplasm, PM, ER, Golgi and tonoplast (Grille et al., 2010; Ramirez-Estrada et al., 2017). Our observations of GFP-TT15 signal in stable Arabidopsis transformants revealed that the TT15 protein is located primarily at the tonoplast or vacuolar membrane. This localization was ascertained by co-localization with the tonoplast markers endosomal-localized Soluble N-ethylmaleimide-sensitive factor Attachment protein Receptor (SNARE) protein VAMP711 and Rab GTPases RabG3c (Rab7D) and RabG3f (Rab7D) previously characterized by Geldner et al. (2009). Rab GTPases and SNAREs ensure membrane-specific tethering and fusion between transport vesicles and target organelles (Uemura and Ueda, 2014). Previously published vacuole proteomes from Arabidopsis cell suspensions (Jaquinod et al., 2007) and rosette leaves (Carter et al., 2004) also mentionned TT15 as being at the tonoplast. Carter et al. (2004) classified it in the “Membrane fusion and remodeling” category. Ramirez-Estrada et al. (2017) demonstrated that TT15 was a peripheral membrane protein, consistent with an absence of transmembrane domain (TMD) in the predicted protein. However they did not detect any consensus amino acid sequences for lipid-mediated reversible post-translational modifications that may be responsible for TT15 transient membrane attachment. Therefore recruitment of TT15 to the tonoplast may rather involve protein-protein interaction or another as yet unidentified mechanism (Ramirez-Estrada et al., 2017).

The tonoplast-located GFP-TT15 fusion protein was also shown to partially colocalize with amyloplasts from the oi2 integument layer. Interestingly, pioneer work with potato tubers reported SG and ASG formation in the amyloplast membrane (Catz et al., 1985). TT15 may be involved in regulating the association of vacuoles and amyloplasts during oi2 cell layer differentiation, similarly to the situation previously described for graviperception in stem and hypocotyl by Saito et al. (2005) and Alvarez et al. (2016), respectively. To our knowledge, such an association has not been reported to date for the mucilage cell layer, but the fact that TT15 is affected in the differentiation of this layer (Berger et al., 2021) suggests that this may be a plausible hypothesis. This mechanism may promote the remobilization of carbohydrates from starch to fuel the biosynthesis of mucilage. Alvarez et al. (2016) propose the formation of a physical tether between vacuole and amyloplast through tonoplast remodeling. It is also possible that the membrane contact site (MCS) may be a zone of exchange for lipids or other molecules between the amyloplast and vacuole membranes, as illustrated with peroxisome by Shai et al. (2016).

Transient expression of a ProCaMV35S-TT15/UGT80B1-YFP fusion infiltrated in *Nicotiana benthamiana* was located both at the PM and in the cytosol by Ramirez-Estrada et al. (2017). On the same line, Pook et al. (2017) located a Pro2xCaMV35S-TT15/UGT80B1-GFP fusion at the PM in stable Arabidopsis transformants. The reason for the discrepancy between our results and these two previous works is unclear. However the fact that in our experimental conditions both GFP-TT15 and TT15-GFP constructs complement the *tt15* mutants and exhibit the same tonoplastic location in accordance with proteomics data from Carter et al. (2004) and Jaquinod et al. (2007) strongly argue towards TT15 being at the tonoplast rather than at the PM. Moreover we did not find any co-localization between TT15-GFP and the PM marker mCherry-NIP1;1. Last but not least, the tonoplastic subcellular localization is relevant with the vacuolar defects observed in *tt15* mutants. We can not rule out the hypothesis that a defect in vacuolar trafficking may affect PM homeostasis (endocytosis, exocytosis) and indirectly explain the cell wall phenotypes observed in *tt15* mutants. An example is provided by the TGN-localized ECHIDNA (ECH) protein which has been shown to be required for the apoplastic secretion of pectin and hemicellulose in oi2 mucilage cells (Gendre et al., 2011; McFarlane et al., 2013) and in the vacuolar sorting pathway for PC accumulation in the endothelium (Ichino et al., 2020). Interestingly, Gendre et al. (2011) reported that mislocalization of the VHA-a1 vacuolar H^+^-ATPase was contributing to *ech* defects at the cell wall. The TGN is highly dynamic and behaves as a central hub for secretion, endocytosis and recycling (Ebine and Ueda, 2015). In absence of TGN-located ECH, cell wall polysaccharides are mistargeted to the vacuole in place of being secreted to the apoplast (McFarlane et al., 2013), which reveals a trafficking connection between the cell wall and the vacuole, at least for polysaccharides. The *tt15/ugt80b1* mutants have been shown to be affected in polysaccharide accumulation in mucilage layer of Arabidopsis seed coat, with *tt15* specifically strengthening primary and secondary cell wall. Moreover, the amount of the major pectin component of the mucilage, rhamnogalacturonan-I (RG-I), was lower in *tt15* as the amount of hemicellulose (galactoglucomannan or GGM) was higher (Zauber et al., 2014; Berger et al., 2021). Collectively these results are compatible with the fact that TT15 is at the tonoplast as most non-cellulosic polysaccharides including hemicelluloses are synthesized in Golgi before being secreted to the apoplast with the contribution of ECH.

### TT15 Contributes to Vacuole Biogenesis and Maintenance

The plant tonoplast has been shown to be organized into microdomains (Ozolina et al., 2013; Yoshida et al., 2013) and to contain SG and ASG (Yoshida and Uemura, 1986; Tavernier et al., 1993; Yamaguchi and Kasamo, 2001). Albeit membrane lipid homeostasis is recognized as being critical in plant vacuolar trafficking, development and response to biotic and abiotic stresses ((Zhang et al., 2015; Sandor et al., 2016; Boutté and Jaillais, 2020), the biological roles of SG and ASG in these processes remain unclear. Here, by monitoring vacuolar deposition of flavanols in seed coat endothelium with vanillin staining, we revealed that a defect in sterol glucosylation due to *TT15* disruption perturbs vacuole biogenesis. Small vacuoles stop increasing in size from the heart stage onwards, suggesting that tonoplast elongation and/or vesicle fusion has been interrupted in *tt15*. The following step is vacuolar degenerescence and necrotic cell death due to release of vacuolar proteases and flavanols in the cytosol. A relationship between vacuole development and SG has previously been observed in fungi, that may put some light on the mechanisms at play in Arabidopsis. Ergosteryl-β-glucoside (EG) is a major class of glycolipids in fungi. Disruption of the steryl-β-glucosidase Egh1 was shown to cause an abnormally fragmented vacuole morphology in *Saccharomyces cerevisiae* (Watanabe et al., 2015). This phenotype was suggested to be due to the accumulation of EG in the vacuole, pointing to a negative role of EG in vacuole fusion, but the precise mechanism involved is unclear (Hurst and Fratti, 2020). Flavonoid metabolism was also shown to affect vacuolar morphology (Abrahams et al., 2003; Baxter et al., 2005; Kitamura et al., 2010; Rosado et al., 2011; Appelhagen et al., 2014), however the molecular triggers still remain to be identified. We can not preclude that the small amount of flavanols that are accumulated in *tt15* endothelial cells interfer with the vacuolar phenotype caused by SG shortage. Investigating further vacuolar morphology in *tt15 tt4* endothelial cells may help answer this question.

We demonstrated in Arabidopsis roots that disruption of the *TT15* gene caused an increase in tonoplast fluidity. Membrane biophysics is characterized by two main parameters: fluidity, as a measure of molecule rotation and diffusion within the membrane; and order, comprising structure, microviscosity and membrane phases (Sandor et al., 2016). SG exhibit the ability to decrease membrane fluidity and to order membranes into microdomains called lipid rafts, defined as detergent-resistant (DRM) or detergent-insoluble (DIM) membranes (Laloi et al., 2007; Grosjean et al., 2015). Previous studies with plant PM have shown that SG and ASG efficiently order membranes and as a consequence reduce their fluidity (Laloi et al., 2007; Halling et al., 2008; Grosjean et al., 2015). Here, we could extend this observation to the tonoplast using a genetic approach based on the characterization of the *tt15-2* mutant affected in SG formation by FRAP. Our study suggests that SG shortage increases membrane fluidity. The biological significance of this observation is important as membrane fluidity was shown to regulate cellular processes such as membrane fusion in yeast (Hurst and Fratti, 2020) or defense signalling in tobacco (Sandor et al., 2016). The modification of tonoplast fluidity is therefore very likely to also affect the function of tonoplastic membrane proteins and membrane remodeling.

### Role of TT15 in Flavanol Trafficking and Vacuolar Sequestration

Flavanol deposition is affected in endothelium, not in micropylar and chalazal cells of *tt15* developing seeds, which suggests that endothelium-specific molecular factors that remain to be determined contribute to this phenotype. Potential candidates which disruption gives a similar seed coat pigmentation pattern than *tt15* are TT1, TT9 and TT16 (Nesi et al., 2002; Sagasser et al., 2002; Ichino et al., 2014). Notably a microscopic investigation of developing seeds revealed that the *tt9* mutant exhibited a similar phenotype as *tt15*. This finding prompted us to analyze the functional relationships between TT9 and TT15. The *tt9* mutant previously isolated by Koornneef (1990) and mapped by Shirley et al. (1995) was shown to be allelic to *gfs9 (green fluorescent seed)* identified as a sorting mutant for vacuolar storage proteins by Fuji et al. (2007). The *tt9/gfs9* mutant exhibits pale seeds, mis-sorting of vacuolar storage proteins, vacuole fragmentation, aggregation of enlarged vesicules, abnormal Golgi morphology and many autophagosome-like structures. AtTT9/GFS9 is localized at the Golgi apparatus, and exhibits strong sequence homology with the *Drosophila melanogaster* Endosomal maturation defective (Ema) protein (Ichino et al., 2014). Ema and its human orthologue C-type LECtin 16A (CLEC16A) cooperatively function with the HOmotypic fusion and Protein Sorting (HOPS) complex and endosomal-localized Soluble N-ethylmaleimide-sensitive factor Attachment protein REceptors (SNARE) proteins to promote lysosomal protein sorting and autophagosome development (Kim et al., 2010; Kim et al., 2012; van Luijn et al., 2015; Pandey et al., 2019). The mechanisms linking endomembrane trafficking to flavanol accumulation in the vacuole (Figure 10A) involve TT9 (Ichino et al., 2014). We discovered that the *tt9-1/gfs9-4* mutant exhibits an endothelium degeneration syndrome that is similar to the one observed in *tt15*. Intriguingly, cell death was suppressed in the double mutant *tt15-2 tt9-1*. Furthermore, vacuole development in ii1 cells exhibited a novel phenotype that did not recapitulate flavanol accumulation observed in wild type. These observations suggest that: 1) if both TT15 and TT9 function in preserving vacuole homeostasis and cell viability, they act in different genetic pathways, since both proteins have a different subcellular localization; 2) the normal development of flavanol-accumulating vacuoles is not essential for cell survival, as shown in the double mutant (Figure 10, B and C). TT9 is required for vacuolar development through vesicle and possibly autophagosome fusion at vacuoles (Ichino et al., 2014). Our results point to TT15 as being important for tonoplast homeostasis, but vesicle fusion at vacuole does not seem to be affected in *tt15*. Therefore both functions are likely to trigger cell death when disrupted, but for different reasons that would complement each other in the double mutant. Apart from a defective accumulation of flavanols in the seed coat, the *tt9* mutants were reported to exhibit vacuolar development and trafficking defects including vacuole fragmentation, mis-sorting of 12S storage proteins and cytoplasmic accumulation of autophagosome-like structures (Shirley et al., 1995; Ichino et al., 2014). In contrast, *tt15* does not exhibit mis-sorting of storage proteins (Ichino et al., 2014) neither vacuole fragmentation (this study) and vacuole development is only partially restored in a double mutant background. Both TT15 and TT9 are peripheral membrane proteins (Ichino et al., 2014; Ramirez-Estrada et al., 2017), meaning that they are likely to move from one subcellular compartment to another.

**Figure 10.**
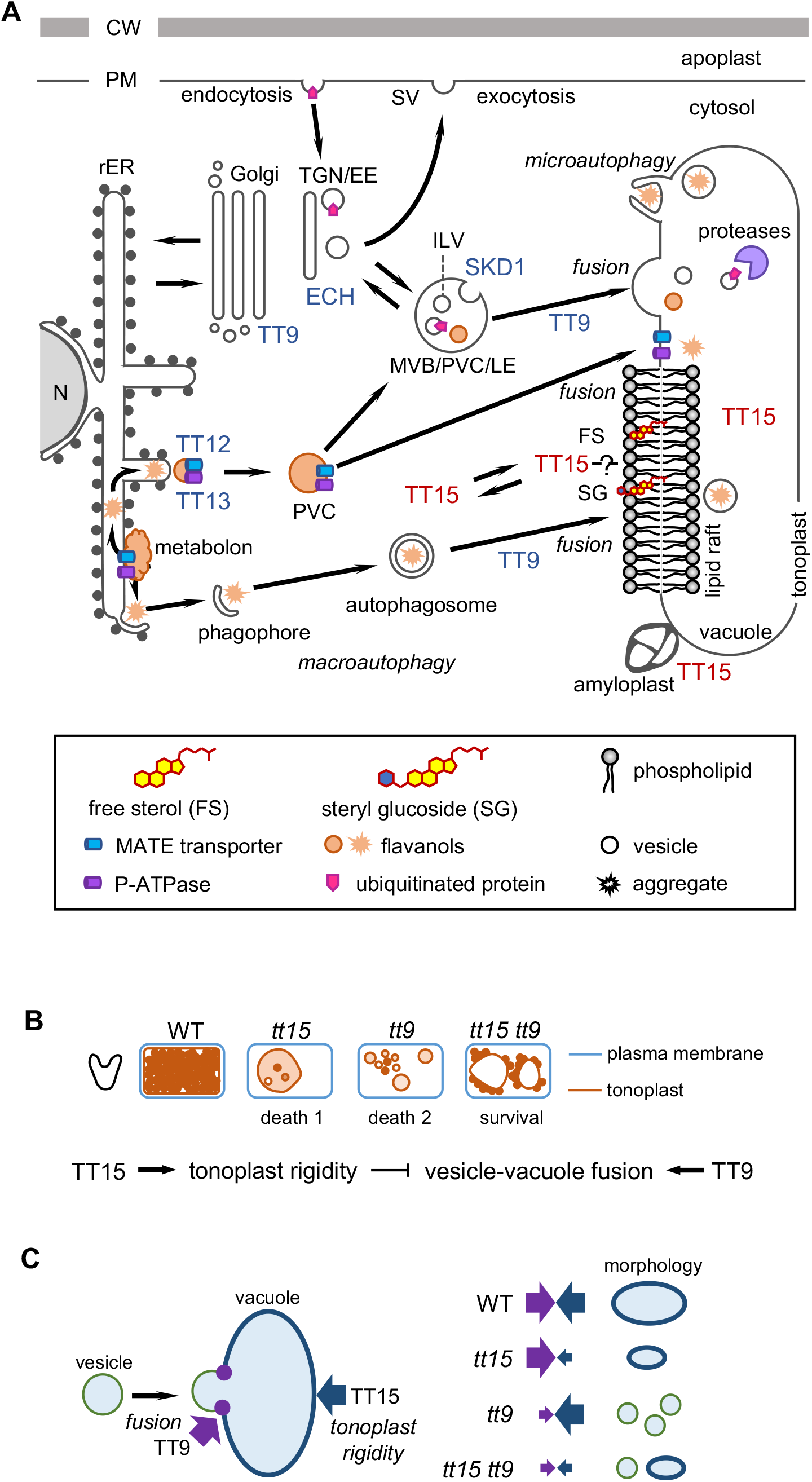
Working Model for the Function of UGT80B1/TT15 in Vacuole Biogenesis and Maintenance at the Seed Coat Endothelium. **(A)** Schematics of the endomembrane trafficking system (adapted from Shimada et al., 2018). TT15 subcellular localizations are shown (in red), together with the ones of other endomembrane-related actors involved in the vacuolar sorting pathway for flavanols (in blue). Vacuolar homeostasis, flavanol deposition, degradation of ubiquitinated proteins and autophagy require TT15 for modulation of tonoplast fluidity by glucosylated sterols concentrated at the level of rafts (microdomains). The presence of TT15 at amyloplast-vacuole contact sites suggests functional relationships between both organelles. In absence of TT15, vacuolar homeostasis (biogenesis and maintenance) is disrupted and tonoplast collapses, enabling vacuolar processing enzymes to degrade the cytosolic machinery which leads to cell death. CW, cell wall; FS, free sterols; ILV, intralumenal vesicle; MVB/PVC/LE, multivesicular body/prevacuolar compartment/late endosome; N, nucleus; PM, plasma membrane; rER, rough endoplasmic reticulum; SG, steryl glucosides; SV, secretory vesicle; TGN/EE, trans-Golgi network/early endosome. **(B)** Genetic relationship between TT15 and TT9. A schematic interpretation of the situation observed in endothelial cells stained with vanillin at the heart stage of embryo development, is proposed. The accumulation of flavanol-containing vesicles in the vacuole lumen is perturbed differently in *tt15* and *tt9*. The defect is partially suppressed in the double mutant, suggesting that other factors involved in this process, beside TT9 and TT15, are impacted by the mutations. Both *tt15* and *tt9* perturb vacuole biogenesis and maintenance, resulting in precocious vacuole collapse and cell death, however through different genetic routes that compensate each other in the double mutant. **(C)** Mechanistic model for the role of TT15 in vacuole dynamics. The increase of tonoplast fluidity caused by *tt15* partially compensates for defective vesicle fusion with the tonoplast due to *tt9*.

The flavonoid phenotypes of *tt15* and *tt9* seeds are similar. Mutant seeds exhibit a strong reduction in total flavanols (soluble and unsoluble flavanols) specifically in endothelial cells, as analyzed with LC-MS and vanillin-based histochemistry (Routaboul et al., 2012; Ichino et al., 2014)(this study). On the other hand, the flavonol fraction remains unchanged apart from quercitrin (quercetin 3-O rhamnose, QR) which is slightly reduced. QR is accumulated essentially in the seed envelopes (Routaboul et al., 2006). The physiological significance of this flavonol phenotype is still unclear. Notably natural variation at the level of the *TT15* locus was observed for both PC and QR seed contents (Routaboul et al., 2012), suggesting an adaptive value for both traits. We can not rule out the hypothesis that flavonoid release from the abnormally developed endothelial vacuoles in *tt15* and *tt9* mutant seed coats triggers cell death. Here, we observed that flavanols (EC and PC) but not anthocyanins or flavonols disrupt cell homeostasis, which may possibly be due to the specific physicochemical properties of decompartimented PC to interact with proteins, carbohydrates and metal ions from the cellular machinery (Hagerman and Butler, 1981; Porter, 1992) and to their cytotoxicity (Dixon and Sarnala, 2020).

The expression of a dominant-negative form of AtSKD1 (SKD1^E232Q^) under the control of the 35S promoter in tobacco induces alterations in the endosomal system leading to cell and plant death (Haas et al., 2007). Interestingly, Arabidopsis plants overexpressing dominant-negative *AtSKD1* constructs under the control of the *GLABRA2* (*GL2*) promoter that restricts expression to trichomes and non-hair cells in the root epidermis, were viable and shown to exhibit a *tt* phenotype and seed mucilage defect (Shahriari et al., 2010a; Shahriari et al., 2010b). Both phenotypes were not characterized further but were supposed to be connected by cell death due to a perturbation in the trafficking of soluble cargo to the vacuole, causing its fragmentation and ultimately cell death (Shahriari et al., 2010b). The same hypothesis may be proposed to explain the *tt15* phenotypes. Thus our results may point to a role played by endosomal and MVB sorting in vacuolar accumulation of PC. On the same line, Gonzalez et al. (2016) showed that TTG2, which disruption causes a *tt* phenotype and an absence of mucilage, controls vacuolar transport of PC by regulating the expression of genes encoding the tonoplastic MATE transporter TT12/DTX41 and P_3A_-ATPase TT13/AHA10. *TT15* regulatory role in the maintenance of endothelial vacuoles (this study) may act at the post-translational level by modulating the activity of either one or both proteins. Relevant with this information, SG have been shown to stimulate the activity of a tonoplastic H^+^-ATPase in rice cell cultures (Yamaguchi and Kasamo, 2002). However *tt15*, *tt12* and *tt13* do not exhibit exactly the same endothelial phenotype, which suggest that other actors may also be involved. As a corollary, we did not observe GFP-TT12 mislocalization in a *tt15* background. Interestingly the pattern of flavanol accumulation and vacuole morphology in *tt15 tt9* endothelium resembles the one observed previously in *tt12* (Debeaujon et al., 2001; Kitamura et al., 2010; Appelhagen et al., 2015) and *tt13* (Appelhagen et al., 2015). In this context we can not rule out the hypothesis that EC transport activity may be impaired in *tt15* indirectly through perturbation of TT9 function as a mediator of vesicle-vacuole fusion.

Altogether, our findings shed light on novel functions of TT15 sterol glucosyltransferase in vacuolar biogenesis and trafficking and their involvement in seed development, germination and flavanol transport to the vacuole. Figure 10 proposes a working model visualizing the respective positions of TT15 and TT9 proteins at the level of the vacuolar trafficking pathways, together with other major actors of the flavanol trafficking pathway. As a basis for future investigations, we speculate that the modulation of tonoplast fluidity through remodeling of its SG composition by TT15 regulates endomembrane-related mechanisms including the fusion of vesicles and autophagosomes to the tonoplast, microautophagy and the activity of tonoplast-localized membrane proteins.

## METHODS

### Plant Materials and Growth Conditions

The Arabidopsis (*Arabidopsis thaliana*) lines used in this study (Supplemental Table S1) were *tt15-1* (Focks et al., 1999; Appelhagen et al., 2014), *tt15-2* (COB16) (Routaboul et al., 2012), *tt15-6* (DNF6) (Nesi, 2001), *tt15-7* (EAL136) (Berger et al., 2021), *tt15-8* (Salk_021175) (Pook et al., 2017), *tt15-9* (Salk_103581) (Stucky et al., 2015), *tt4-8* (DFW34) (Debeaujon et al., 2003) and *tt9-1/gfs9-4* (Koornneef, 1990; Shirley et al., 1995; Ichino et al., 2014). The *tt15-2*, *tt15-6*, *tt15-7 and tt4-8* alleles are in wild-type Wassilewskija (Ws-4) accession and were obtained from the INRAE Versailles T-DNA collection (Brunaud et al., 2002). The *tt15-8* and *tt15-9* alleles in wild-type Columbia (Col-0) accession are from the Salk Institute T-DNA collection (Alonso et al., 2003) and were obtained from the NASC stock center. The *tt15-1* allele in Col-2 background and the *tt9-1* allele in L*er* background were provided by Christophe Benning and Maarten Koornneef, respectively. The double mutant between *tt15-2* and *tt9-1* was obtained by crossing, using *tt15-2* as female. F2 plantlets were genotyped for *tt15-2* as described in Berger et al. (2021) and for *tt9-1/gfs9-4* by sequencing after PCR amplification with TT9-F3 and TT9-GST3 primers (Supplemental Table S5). The double mutant with *tt4-8* was selected on the basis of its pale yellow seed colour and absence of anthocyanins in vegetative parts, after checking for the presence of *tt15-2* as described above. For reciprocal crosses, F1 hybrid seeds produced by WT mother plants with a *tt15* pollen donor are referred to as WT x *tt15* F1 seeds and those from *tt15* mother plants with a WT pollen donor as *tt15* x WT seeds. For FRAP experiments, the transfer of GFP-NRT2-7 in a *tt15-2* background was done by crossing *tt15-2* with a transformant obtained from Chopin et al. (2007) expressing the cassette in Ws-4 background. For co-localization experiments, fluorescent subcellular compartment markers (Supplemental Table S4) were crossed with a representative transformant expressing GFP-TT15.

For *in vitro* cultures, seeds were surface-sterilized, sown in Petri dishes containing Gamborg B5 medium (Duchefa, The Netherlands) supplemented with 3% sucrose and 0.8% agar, stratified for 3 days at 4°C in the dark and grown for 3 to 12 days at 20°C with a 16-h light/8-h dark cycle and 70% relative humidity. Plant growth in glasshouse was realized on compost (Tref BV, The Netherlands) fertilized with Plant-Prod nutritive solution (Fertil, France) at around 23°C / 15°C (day/night). Observation of seed lethality was performed on plants grown in controlled conditions in a growth chamber settled at 21°C/18°C (day/night), 16h lighting and 65% relative humidity.

### Seed Germination Assays

Seed lots from WT (Ws-4, Col-0) and *tt15* mutants to be compared were obtained from plants grown at the same time and in the same environmental conditions. Seeds from a bulk of four plants from each genotype was sown in triplicate in 6-cm Petri dishes (50-60 seeds per dish) on 0.5% (w/v) agarose supplemented with ABA (Junda Pharm Chem Plant Co., China) or the gibberellin biosynthesis inhibitor paclobutrazol (Syngenta, Switzerland). Seeds were stratified at 4°C for three days in the dark and transferred in a growth cabinet (continuous light, 25°C, 70% relative humidity). Germination (emerged radicle) and/or cotyledon greening were scored four days after transfer to light.

### Constructs

All primers used for plasmid constructs are listed in Supplemental Table S5. For construction of the *ProTT15:uidA* transgene, a region located **-**2003 to **-**1 bp relative to the *TT15* translational start codon was amplified with the ProTT15-SalI and ProTT15-SmaI primers using a proof-reading Taq polymerase (Phusion; Thermofisher, USA) and cloned in TOPO vector (Thermofisher, USA). After validation by sequencing, *ProTT15* was digested by SalI-SmaI and cloned at the SalI-SmaI sites of the pBI101 binary vector (Clontech, USA) for plant transformation. To construct the GFP-TT15 translational fusion protein, the *TT15* coding sequence (CDS) was amplified from clone U22595 (SSP pUni clone BT005834 from the Salk Institute, USA) with TT15-ATG-attB1 and TT15-END-attB2 primers using Phusion Taq polymerase. The PCR product was then cloned into pDONR207 by BP recombination (Gateway BP Clonase enzyme mix, Invitrogen, USA). After validation by sequencing, TT15 CDS was transferred into the binary vectors pMDC43 and pMDC83 (Curtis and Grossniklaus, 2003) by LR recombination. Promoter activity and subcellular localization of translational fusions with GFP were investigated after stable transformation of Arabidopsis plants according to the floral dip method (Clough and Bent, 1998). Around fifteen independent transformants per construct were obtained, among which two were selected for further characterization on the basis of their representative behaviour.

### Light Microscopy

Observations were performed with an epifluorescence microscope (Zeiss Axioplan 2, Germany) equipped with Nomarski differential interference contrast optics. Silique clearing for quantification of seed lethality was performed as described by Surpin et al. (2003), on 25 siliques per genotype (five siliques from five plants) harvested 15 days after flowering. Fixation, resin embedding, sectioning and toluidine Blue O (TBO) staining of seed material were realized as described in Debeaujon et al. (2003). TBO stains polyphenolic compounds (procyanidins and lignins) in greenish-blue, pectin in pink and nucleic acids and proteins in purple. Histochemical detection of GUS activity was performed as described in Debeaujon et al. (2003), using 2.5 mM potassium ferricyanide and 2.5 mM potassium ferrocyanide. Developing seeds were cleared by overnight incubation in a chloral hydrate:glycerol:water (8:1:2, w:v:v) solution. The vanillin assay was used for specific staining of colorless flavanols in bright red in developing seed coat, as described by Debeaujon et al. (2000). ROS (H_2_O_2_) detection was realized according to Bailly and Kranner (2011) using 5-(and-6)-chloromethyl-2’, 7’-dichlorofluorescein diacetate (DCFH-DA; Sigma, USA). Developing seeds were incubated for 15 min in 100 μM DCFH-DA in 20-mM potassium phosphate buffer pH 6.0 at 20°C, and rinsed three times with buffer for 5-min each before observation under UV light. For callose detection in developing seeds, young siliques (2-3 dap) were fixed, hydrated, softened and stained in fresh 0.1% Aniline Blue (Acros Organics, Belgium) aqueous solution according to Huck et al. (2003). Seed whole mounts were observed under UV light.

### Confocal Laser Scanning Microscopy

Observations were performed using a Leica SP5 or a Leica SP8 spectral confocal laser scanning microscope (Leica Microsystems, Germany) fitted with 20x or 63x water-immersion objectives. A 488-nm line from an argon laser was used to excite GFP, and fluorescence was detected in the 501- to 598-nm range. A 561-nm line from a He-Ne laser was used to excite mCherry, and fluorescence was detected in the 571- to 668-nm range. Chlorophyll autofluorescence was detected in the 615- to 715-nm range. Images were false-colored in green (GFP) or magenta (mCherry or chlorophyll autofluorescence) with the ImageJ software (Schneider et al., 2012) and processed using the FigureJ plugin (Mutterer and Zinck, 2013). The fluorescent pH indicator BCECF-AM (Life Technologies, USA) was used to stain the acidic lumen of the vacuoles according to Scheuring et al. (2015). Seedling roots were incubated in a 10-μM BCECF solution during 1 h in the dark and rinsed before confocal imaging. The wavelengths for excitation and emission were 488 and 520 nm, respectively.

Vacuolar membrane fluidity was determined by FRAP using the NRT2.7 nitrate transporter fused to GFP (Chopin et al., 2007) as a tonoplastic intrinsic marker in Ws-4 and *tt15-2* backgrounds. Images were acquired on epidermal cells of the elongation zone of 4-day-old seedling roots. Measurements were performed using the FRAP wizard of Leica SP5 microscope with a 63x objective. Rectangular Regions of Interest (ROI) of 8.9 μm^2^ were designed. Fluorescence intensity data were collected from bleached area (ROI1), total fluorescence area (ROI2) and background area (ROI3). Pre-bleaching and post-bleaching imaging was performed with the 488-nm line of an Argon laser at 100% output and 10% transmission. For bleaching, one scan of ROI1 was done at 100% transmission of 405-nm diode and 458-nm, 476-nm and 488-nm lines of Argon laser with “zoom in” method of bleach. Scans were done with a minimized time frame of 0.189s. Time course of acquisition was as follows: pre-bleach: 10 frames, bleach: 1 frame and post-bleach: 200 frames. Fluorescence intensity data were normalized with the easyFRAP software (Rapsomaniki et al., 2012) using full-scale normalisation method. T_half_ (half maximal recovery time) and mobile fraction were computed after curve fitting using single term equation. Wilcoxon test was used to calculate the statistical significance of T_half_ and mobile fraction results between Ws-4 wild type and *tt15-2* mutant (significance at P<0,001).

### Accession numbers

Sequence data from this article can be found in the Arabidopsis Genome Initiative (TAIR) or EMBL/GenBank data libraries under the following accession numbers: TT15/UGT80B1 (AT1G43620, NP_175027), UGT80A2 (AT3G07020, NP_566297), TT4 (AT5G13930, NP_196897), TT9 (AT3G28430, BAH57204).

## SUPPLEMENTAL DATA

**Supplemental Figure S1.** Biosynthesis of steryl glucosides and related end-products.

**Supplemental Figure S2.** Biosynthesis of flavonoids and related end-products.

**Supplemental Figure S3.** *TT15* mutant alleles used in this study.

**Supplemental Figure S4.** *transparent testa* mutants with a similar pattern of seed coat pigmentation than *tt15* also exhibit some seed lethality (Supports Figure 1).

**Supplemental Figure S5.** The *tt15* mutant alleles produce smaller seeds than wild types (Supports Figure 1).

**Supplemental Figure S6.** Phenotypic diversity of *tt15* seedlings (Supports Figure 1).

**Supplemental Figure S7.** Impact of the *tt15* mutations on seed physiology (Supports Figure 2).

**Supplemental Figure S8.** The *tt15* mutant seed developmental phenotypes in Col-0 background are similar to the ones in Ws-4 (Supports Figure 3).

**Supplemental Figure S9.** Flavanol depletion suppresses endothelium browning and breakdown (Supports Figure 3).

**Supplemental Figure S10.** The *TT15* promoter is active in reproductive and vegetative organs (Supports Figure 5).

**Supplemental Figure S11.** *TT15* is expressed in vegetative and reproductive organs similarly in Ws-4 and Col-0 accessions (Supports Figure 5).

**Supplemental Figure S12.** *TT15 in silico* expression data (Supports Figure 5).

**Supplemental Figure S13.** The translational fusions of TT15 with the fluorescent Protein GFP used for subcellular localization studies complement the *tt15* mutant (Supports Figure 6).

**Supplemental Figure S14.** Subcellular localization of GFP Fusions with TT15 in seed coat and root (Supports Figure 6).

**Supplemental Figure S15.** TT15 does not co-localize with Golgi, TGN, late endosome and plasma membrane markers (Supports Figure 6).

**Supplemental Figure S16.** TT15 absence does not significantly impact flavonoid gene expression in seeds.

**Supplemental Figure S17.** Vacuole morphology is not affected in absence of TT15 in roots (Supports Figure 8).

**Supplemental File S1.** Supplemental tables 1 to 5. Mutants **(S1)**; Flavonoids **(S2)**; Seed lethality **(S3)**; Fluorescent markers **(S4)**; Primers **(S5)**.

**Supplemental Movie 1** (Supports Figure 5). Z-Stack of GFP-TT15 in a cotyledon from a 7-day-old seedling, focusing on a stomata (spinning-disk; 67 frames).

**Supplemental Movie 2** (Supports Figure 5). Dynamics of GFP-TT15 in a root tip of a 3-day-old Arabidopsis seedling, from the meristematic zone to the elongation zone (time series 20 planes in 3 s).

**Supplemental Movie 3** (Supports Figure 5). Dynamics of GFP-TT15 in a root tip of a 7-day-old Arabidopsis seedling, at the elongation zone (spinning-disk; time series 167 planes in 24 s).

**Supplemental Movie 4** (Supports Figure 5). Dynamics of GFP-TT15 and mCherry-GOT1p in a root tip of a 3-day-old Arabidopsis seedling, at the elongation zone (time series 25 planes in 4 s).

## ACKNOWLEDGMENTS

We are very grateful to Maarten Koornneef for providing *tt9-1* seeds, Christoph Benning for *tt15-1* seeds, and Sylvie Ferrario-Méry for seeds of the GFP-NRT2.7 line. We thank Lucille Pourcel, Guillaume De Lagarde and Nathan Leborgne for contribution to the experimental work. This work was funded by the European Commission (FOOD-CT-2004-513960 “FLAVO” and FP7 Environment Grant Award Number 311840 “EcoSeed”) and has benefited from the support of IJPB’s Plant Observatory technological platforms. The IJPB benefits from the support of Saclay Plant Sciences-SPS (ANR-17-EUR-0007).

## AUTHOR CONTRIBUTIONS

E.A., A.B., F.P., A.F., A.T., S.C., H.S., S.V., O.G., N.N. and I.D. performed research and analyzed data; E.A., L.L., A.M.P. and I.D. designed research; E.A. and I.D. wrote the paper.

